# IDPForge: Deep Learning of Proteins with Global and Local Regions of Disorder

**DOI:** 10.64898/2026.03.25.714313

**Authors:** Stefano De Castro, Oufan Zhang, Zi Hao Liu, Julie D Forman-Kay, Teresa Head-Gordon

## Abstract

Although machine learning has transformed protein structure prediction of folded protein ground states with remarkable accuracy, intrinsically disordered proteins and regions (IDPs/IDRs) are defined by diverse and dynamical structural ensembles that are predicted with low confidence by algorithms such as AlphaFold and RoseTTAFold. We present a new machine learning method, IDPForge (**I**ntrinsically **D**isordered **P**rotein, **FO**lded and disordered **R**egion **GE**nerator), that exploits a transformer protein language diffusion model to create all-atom IDP ensembles and IDR disordered ensembles that maintains the folded domains. IDPForge does not require sequence-specific training, back transformations from coarse-grained representations, nor ensemble reweighting, as in general the created IDP/IDR conformational ensembles show good agreement with bot NMR and SAXS solution experimental data, and options for biasing with experimental restraints are provided if desired. We envision that IDPForge with these diverse capabilities will facilitate integrative and structural studies for proteins that contain intrinsic disorder, and is available as an open source resource for general use.

## 1 Introduction

Structures of biomolecules have driven functional insight ever since Watson and Crick solved the structure of DNA, establishing the structure-function paradigm that has been critical to progress in molecular biology and biochemistry. X-ray crystallography and cryo-EM along with nuclear magnetic resonance (NMR) have informed the traditional structure-function paradigm for folded proteins for many decades. The recent advent of machine learning (ML) models for protein structure prediction, most notably AlphaFold (AF)^1,2^ and RoseTTAFold ^3^, have made stunning breakthroughs in producing accurate ground state structures of monomeric folded proteins of high quality, similar to that of experimental structures. Although static structures of folded proteins can yield significant insights, protein function is inherently controlled by dynamics. ^4–8^ But the surprising depth of deep learning models suggest that they too can create not just a single structure, but a diverse and representative protein structure-dynamical space for functional insight.

In fact the ML/AI field has seen an emergence of a number of approaches to learn structural variants of folded states. ^9^ One expected class of ML algorithms are generative models that learn conformational distributions from molecular dynamics (MD) data via autoencoders, Boltzmann generators^10^ and flow-matching^11^, or to drive enhanced conformational sampling ^12,13^ in learned lower manifolds of collective variables^14^. Notably, denoising diffusion probabilistic models (DDPMs) have emerged as a powerful framework for structural variant sampling and generative protein design ^2,15–20^. Compared to traditional generative approaches, DDPMs provide a more flexible mechanism to incorporate diverse conditioning signals, such as functional constraints, secondary structure preferences, or binding site requirements, enabling the generation of proteins tailored to specific tasks^15–17,21^. Other approaches involve inference time manipulations of structure prediction models. The Meiler lab cleverly demonstrated that reducing the depth of the input multiple sequence alignments (MSAs) in AF2 led to predictions of protein structures in multiple conformational states ^22^, an idea that was later elaborated upon by other groups using sequence clustering or sub-sampling the MSAs to isolate the origin of conformational states of certain classes of folded proteins. ^23,24^

But approximately two-thirds of the canonical human proteome are predicted to be within intrinsically disordered proteins (IDPs) or intrinsically disordered regions (IDRs) that do not adopt a dominant folded structure, but instead utilize their conformational range to execute diverse cellular functions. ^25,26^ Because X-ray crystallography and cryo-EM methods are non-viable for this class of protein, computational models must bridge the gap by creating IDP ensembles that agree with the highly averaged experimental solution data from NMR, small angle X-ray scattering (SAXS), single molecule Förster Resonance Energy Transfer microscopy (smFRET) and pulsed EPR double electron-electron resonance (DEER-EPR). ^27–29^ The generation of putative disordered ensembles can be based on knowledge based methods such as IDPConformer Generator ^30^, atomistic molecular dynamics^31^ or a *C_α_* based coarse-grained representation ^32,33^. These structural “pools” of disordered protein conformers are often reweighted to give an ensemble average that agrees with experimental observables ^34–38^, although there is a risk that the underlying pool is incomplete and missing important populations. Recently generative algorithms have become popular for generating all-atom or coarse-grained IDP ensembles ^39–43^. For example idpGAN uses a generative adversarial network that learns a *C_α_* based coarse-grained representation of disordered sequences, then superseded by a latent diffusion model with enhanced transferability to their test sequences ^41,42^. IDPFold developed by Zhu *et al.* employed a two-stage training of a *C_α_* backbone denoising model with PDB and all-atom molecular dynamics (MD) simulation structures, which showed promising performances on sampling structured and disordered proteins ^43^. Still they often require postprocessing by some type of reweighting step to gain better experimental relevance. Our group introduced a generative recurrent neural network (GRNN), DynamICE (**Dynam**ic **I**DP **C**reator with **E**xperimental Restraints) ^44^, that uses experimental data to bias the torsion angle distributions of an underlying structural pool to generate new conformational states that conform to experimental averages. The significance of the DynamICE approach is that we remove the iterative guesswork about relevant sub-populations and subsequent reweighting of arbitrarily drawn conformations. But the underlying long short term memory (LSTM) architecture uses an internal coordinate representation of protein conformers, which is sub-optimal for meeting distance restraints such as nuclear overhauser effect (NOEs) and paramagnetic relaxation enhancement (PREs). Hence a method that can exploit Cartesian coordinates would be a more natural framework. Furthermore DynamICE like other Bayesian and generative models mentioned above also requires training each new IDP sequence and/or with its own experimental structural restraints, which reduces its generalizability.

Here we shed the limitations of generic generative models and sequence specific training to create experimentally valid conformational ensembles for IDPs as well as IDRs from proteins containing folded domains, with all-atom resolution. We adapt the attention and structure modules from the ESMFold network^45^ to a generative model for disordered protein structures in a Denoising diffusion probabilistic models (DDPM) framework, by supplying an additional noisy structure input as exemplified in RFdiffusion ^16^ and AlphaFlow (ESMFlow) ^11^. Re-purposing established deep learning models has several benefits: it saves the considerable effort in developing a model from scratch that combines sequence and inter-residue feature extraction for polymer-like biomolecules that is already handled well by other methods. Instead we enable structure predictions of disordered regions in conjunction with folded domains with simple adjustments of diffusion steps at inference time. The resulting IDPForge (**I**ntrinsically **D**isordered **P**rotein, **FO**lded and disordered **R**egion **GE**nerator) unites ensemble generation of IDPs and IDRs with folded domains within one model, demonstrating excellent performance in sampling all-atom conformational ensembles in agreement with a broader range of solution experimental data including chemical shifts, J-couplings, NOEs, and PREs from NMR, and radius of gyration from small angle X-ray scattering (SAXS). We also introduce a sampling procedure for guidance based on experimental data types to further align ensemble generation to the experimental averages that are effective for distance restraints such as NOEs and PREs, without additional training cost. With these diverse capabilities we expect that IDPForge can be applied across structural and integrative biology tasks for proteins that contain intrinsic disorder, and is available as an open source resource.

## 2 Results

The general framework of IDPForge is shown in Figure 1, and details of the architecture and training protocols can be found in Methods. We first demonstrate the ability of IDPForge to generate whole single chain IDPs (not on fragments) on a collection of 32 test sequences ^32,41,43^ with available NMR and/or SAXS data (Supplementary Table S1). We also compare IDPForge performance with other published methods, including an all-atom force field a99SB-disp ^31^ and the coarse-grained force field CALVADOS ^32^ and ML models idpGAN^41^, STARLING ^46^, IDPFold ^43^ and later with AFflecto ^47^. In addition to the generation of single chain IDPs, we showcase two inference time generation examples with shifted distributions: 1) sampling with experimental data guidance and 2) sampling local IDRs within the context of folded domains. In the latter case, some proteins in the test set were decided upon based on availability of CS data in order to validate and compare methods. Basic structural validation for bond, angle and torsion statistics of IDPForge over sampled conformations is provided in Supplementary Fig. 1.

**Figure 1:**
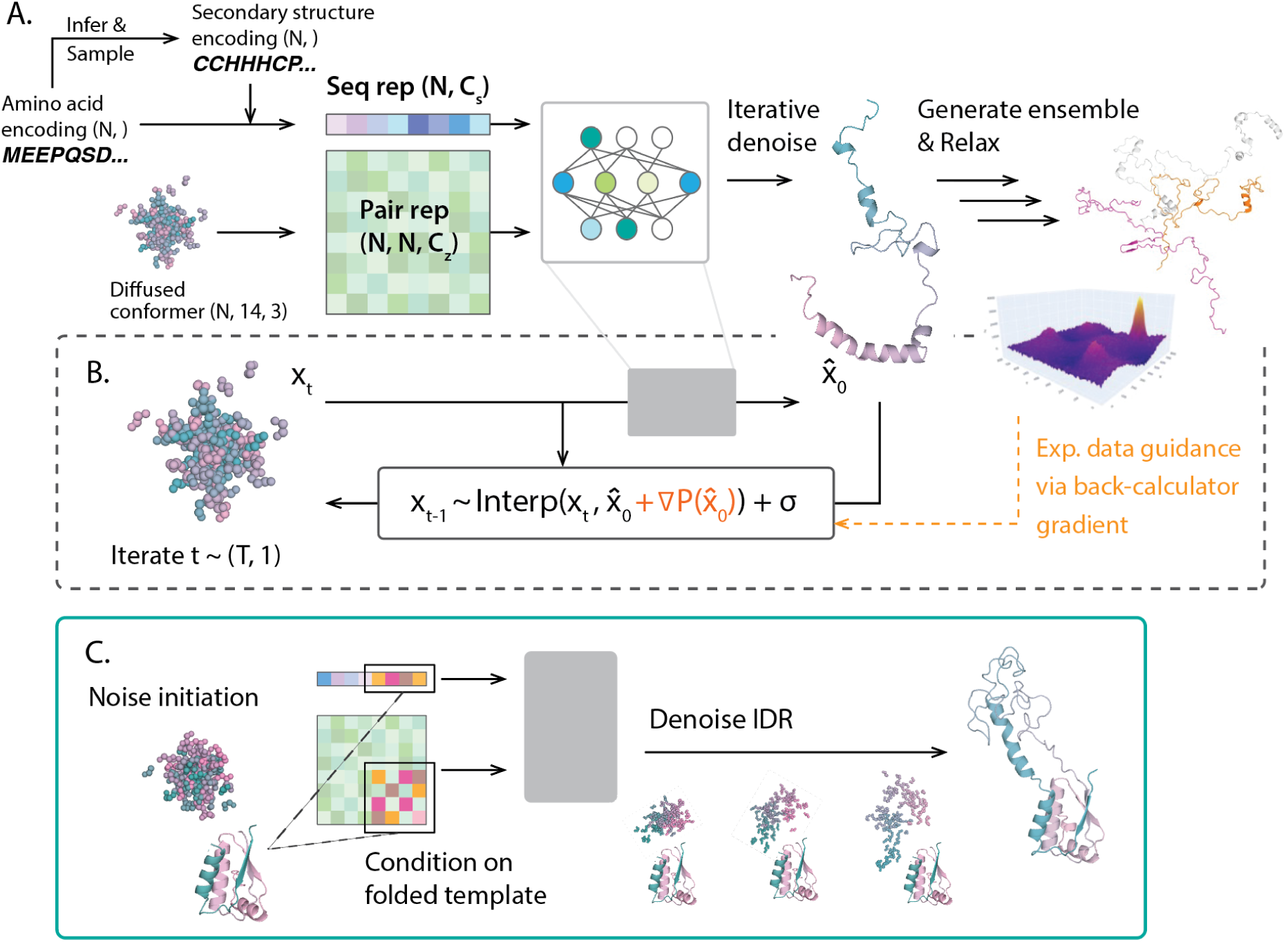
The schematic of IDPForge framework for IDP/IDR prediction. (A) DDPM network that takes a protein sequence and diffused conformer coordinates *x_t_* that are featured into a sequence and a pair representation to predict protein structure *x*_0_ by iterative denoising for time steps *t* = *T*. Secondary structure propensities are inferred from the protein sequence, and candidate secondary structure encodings are sampled based on these propensities using DSSP ^48^ for helices (H) and sheets (E) assignment and further classifies DSSP coils into poly-proline II (P) and other Ramachandran regions as defined in Supplementary Fig. S2. Details of network architecture and input features are discussed in Section 4.5 and Supplementary Section 1.1. Generated structures are relaxed using an empirical force field. Notation: *N* residue number, *C_s_* sequence representation dimension and *C_z_* pair representation dimension. (B) Breakdown of a single denoising step *x_t−_*_1_ → *x_t_* into a network forward pass *x* ^_0_ → *ɛ*(*x_t_*) and a reverse step mapping *x_t−_*_1_ → Interp(*x_t_, x* ^_0_) + *σ* based on Algorithm S2, with optional experimental data guidance via gradients formulated based on back-calculators ∇*P* (*x^* _0_). During sampling, *x_T_* is initiated as random noise and the denoising step is repeated while decrementing the time step *t*. (C) Sampling of IDRs with folded domains via conditioning on a given folded template and partial denoising of the disordered sequence at inference time.

### 2.1 Generating single chain disordered ensembles

Table 1 reports the test set evaluation of 32 generated single chain IDP ensembles across different experimental data types for the various ensemble generation methods. We note that not all 32 IDP proteins have a complete set of all experimental data types; we consider chemical shift (CS) *χ*^2^ performance for 15 sequences with CS values taken from the BMRB^49^, normalized radius of gyration error with respect to reported experiments for 30 IDPs, and 5 proteins that have J-couplings (JC) and NOEs or PREs. We rank performance on methods using the X-EISD Bayesian score ^38^ (see Eqn. S11) for the NMR experimental data types as it takes into account the various sources of experimental and back-calculation uncertainties when measuring the fit between the structural ensemble and the experimental observable ^34,38^; a higher X-EISD score corresponds to a higher likelihood that the structural ensemble is in agreement with the given data type or all data types together. Because the X-EISD scoring function already accounts for experimental and back-calculation error, then the normalized X-EISD score is a meaningful ranking among methods. We also note that although IDPForge is evaluated on 32 test set proteins that does not have above 30% overlap with the training set proteins, all other methods besides IDPConformerGenerator have seen some of the test proteins in their training data set. For example, 13 of the 32 test set proteins were part of the training data for CALVADOS; 9 of the 32 were in the training data for IDPFold; a different set of 13 proteins from the test set were part of the training data for IDPGan; STARLING did not report its training and test set based on protein ID so it is unclear what the overlap is with the test set here (although we expect it to be a large fraction). Nonetheless we believe these comparisons are still useful as performance is measured on a wider range of experimental NMR data on complete IDP sequences and beyond just Rg from SAXS data alone.

**Table 1:** Experimental benchmark of the ensembles generated by IDPForge and other methods. The experimental data for X-EISD calculations include J-Couplings (JC), nuclear overhauser effect (NOE), paramagnetic relaxation enhancement (PRE) and chemical shifts (CS). The X-EISD scores (Eq. S10) are normalized for each protein between 0-1 using Eqn. S11, and consider proteins with available data types reported in Table S1. X-EISD accounts for error and uncertainty for both experiments and back-calculations. The uncertainties in values are reported in the last significant figure. The radius of gyration error with respect to the experimental values Δ*R_g_* are calculated using Eqn. S5. The relative performance among the methods is broken down by test evaluation category in Supplementary Fig. S2.

| | Normalized X-EISD score ↑ | | | | $\chi^2_{CS} \downarrow$ | $\%\vert \Delta R_g\vert /R_g^{exp} \downarrow$ |
| --- | --- | --- | --- | --- | --- | --- |
|  | Total | CS | JC | NOE/PRE |  |  |
| IDPForge | 0.89(0) | 0.91(7) | 0.95(6) | 0.72(8) | 0.15(6) | 6.9(2) |
| IDPConformerGenerator | 0.57(6) | 0.47(1) | 0.82(9) | 0.47(7) | 0.27(2) | 7.4(3) |
| IDPFold | 0.58(2) | 0.62(7) | 0.39(3) | 0.45(6) | 0.27(1) | 11.7(5) |
| idpGAN | 0.20(9) | 0.16(7) | 0.34(7) | 0.72(7) | 0.36(7) | 10.3(2) |
| STARLING | 0.39(5) | 0.47(5) | 0.38(0) | 0.48(9) | 0.20(7) | 4.7(4) |
| CALVADOS | 0.58(4) | 0.67(7) | 0.69(4) | 0.49(1) | 0.27(2) | 5.0(3) |

When averaged across IDPs, IDPForge demonstrates excellent agreement with the NMR solution experimental data, outperforming all other methods when accounting for chemical shifts, J-couplings, and NOEs/PREs as evaluated from the total X-EISD score in Table 1 and Supplementary Fig. S3(A). To better analyze statistical differences in ranking, we also break down the NMR data type score for each protein for each method in Supplementary Fig. S4. While the relative ranking by protein can change, aggregated across the test proteins IDPForge does better on NMR solution data. We find that the generated ensembles by all methods yield *χ*^2^ values within the uncertainties of the CS back calculator (UCBShift ^50,51^) for all IDP proteins as seen in Table 1 and Supplementary Fig. S3(B), with the exception of PaaA2 and the unfolded state of drkN-SH3 which degrades the performance of methods such as idpGAN, IDPFold, and CALVADOS.

Supplementary Fig. S5 sheds light on the *C_α_* CS inconsistencies of these methods from the perspective of secondary structure features derived from NMR versus their heavier weighting of Rg from SAXS data. As already reported for the conformational ensemble derived from SAXS and NMR chemical shift, PaaA2 exists in solution with two preformed helices, connected by a flexible linker, believed to be two types of molecular recognition elements for toxin inhibition. ^52^ Sterckx and co-workers showed that a random coil pool containing no *α*-helices and a pool with *α*-helical populations are both in agreement with SAXS (i.e. Rg), but only ensembles selected from the pool containing the preformed *α*-helices can adequately reproduce the NMR data in addition. Hence the two coarse-grained methods CALVADOS and idpGAN report a higher *χ*^2^ due to their lack of secondary structure features. The observed connection between chemical shifts and secondary structure profile highlights that IDPs are not featureless coil-like structures, and thus may miss an important factor in biological relevance. The drkN-SH3 ensemble generated by IDPFold features a considerable *β*-sheet population between residue 35-50, likely reminiscent of its folded states rather than the unfolded states ^53^, thus leading to its poor fit to the experimental chemical shifts compared to the other methods. IDPForge improves upon chemical shift predictions due to the presence of localized regions of transient helical/turn secondary structure across the sequences of the unfolded state of drkN-SH3.

In regards the *R_g_* error for the 30 test IDPs in Table 1 and Supplementary Fig. 3C, it is found that STARLING and CALVADOS perform the best since these methods emphasize SAXS data in their development, followed by IDPForge and IDPConformerGenerator, while IDPFold and idpGAN perform the most poorly. At the same time the best performing STARLING and CALVADOS methods for *R_g_*perform much worse than other methods on the NMR data. To gain more insight into this imbalance across what should be related data types, i.e. distance restraints from NMR should be commensurate with Rg from SAXS, we compare the subset of 9 disordered proteins for which long MD simulation trajectories using the a99SB-disp forcefield are available^31^ in Table 2. On this test set we see that the a99SB-disp ensembles have the smallest *R_g_* errors while also showing excellent agreement with CS, JC, and NOEs/PREs, indicating that these data types are mutually reinforcing such that the underlying a99SB-disp ensembles are more self-consistent. Thus it appears that STARLING and CALVADOS are mostly designed to create ensembles that match experimental *R_g_*’s, but are less consistent with the other NMR data types. The strength of IDPForge is its overall agreement and uniformity in performance across local (CS and JC) and global (NOE/PRE and *R_g_*) data types.

**Table 2:** Experimental benchmark of the ensembles generated for test proteins compared to molecular dynamics. The subset of proteins for which MD using the a99SB-disp forcefield are available ^31^: A*β*40, ACTR, Ash1, *α*-Syn, drkN-SH3, *N_TAIL_*, *p*15*^P AF^*, PaaA2 and Sic1. The experimental data for X-EISD calculations include J-Couplings (JC), nuclear overhauser effect (NOE), paramagnetic relaxation enhancement (PRE) and chemical shifts (CS). The X-EISD scores (Eq. S10) are normalized for each protein between 0-1 using Eqn. S11, and consider proteins with available data types reported in Table S1. X-EISD accounts for error and uncertainty for both experiments and back-calculations. The uncertainties in values are reported in the last significant figure. The *R_g_* error with respect to the experimental values Δ*R_g_* are calculated using Eqn. S5.

| | Normalized X-EISD score ↑ | | | | $\chi^2_{CS} \downarrow$ | $\%\vert \Delta R_g\vert /R_g^{exp} \downarrow$ |
| --- | --- | --- | --- | --- | --- | --- |
|  | Total | CS | JC | NOE/PRE |  |  |
| IDPForge | 0.78(2) | 0.89(2) | 0.90(4) | 0.69(9) | 0.13(5) | 5.9(7) |
| IDPConformerGenerator | 0.62(7) | 0.50(8) | 0.91(0) | 0.56(6) | 0.26(2) | 5.3(5) |
| IDPFold | 0.48(2) | 0.50(5) | 0.28(2) | 0.43(5) | 0.28(5) | 7.5(3) |
| idpGAN | 0.26(0) | 0.13(9) | 0.55(4) | 0.73(0) | 0.37(6) | 4.7(8) |
| STARLING | 0.21(0) | 0.28(8) | 0.19(9) | 0.38(1) | 0.22(9) | 4.7(7) |
| CALVADOS | 0.51(9) | 0.58(4) | 0.72(4) | 0.42(9) | 0.30(0) | 4.8(3) |
| a99SB-disp | 0.77(2) | 0.84(3) | 0.52(9) | 0.87(1) | 0.21(5) | 2.9(1) |

### 2.2 Generation of IDP/IDR ensembles guided by experimental data

For cases in which experimental information is available across data types, using it to bias ensemble generation has proved fruitful for IDP structural ensemble analysis. ^28^ For example, some of the IDP test proteins such as drkN-SH3 and RNaseA are folded proteins, but the experimental data that is available relates to the unfolded state. Experimental data also contains environmental signatures, for example that *α*-Syn is disordered in aqueous solution but is folded into helices when membrane-bound. We also showed in our DynamICE work that Hst5 structural ensembles depend on solvent conditions, which is also reflected in J-couplings. ^44^ So while IDPForge predicts disordered states first, it can be sculpted by experimental data that changes secondary structure (influenced by J-couplings), or long-range interactions (like NOEs/PREs) or collapsed and expanded states (such as Rg) dependent on environment or post-translational modifications (PTMs).

Hence we augment IDPForge with sampling biases to further align the generated ensembles towards better agreement with the experimental data as shown in Table 3. We highlight that this sampling guidance strategy requires no training but is done at the inference stage. When comparing the X-EISD score for each IDP (Supplementary Fig. S3(A) and S4), IDPForge ranks at or near the top by total X-EISD score across all proteins with the exception of *α*-Syn, whose disordered state is a precursor to toxic fibers found in Parkinson’s disease^54,55^, and the intrinsically disordered cyclin-dependent kinase (CDK) inhibitor Sic1^56^. In both cases this arises due to PRE error, a problem shared by the other methods including for CALVADOS, STARLING, and a99SB-disp simulations. Table 3 shows that IDPForge when guided with long-range PRE experimental restraints (only using PREs for residue number differences greater than 10) significantly reduces the mean absolute error (MAE) error for this property for both Sic1 and *α*-Syn, while maintaining good agreement for other data types. This IDPForge capability allows ensemble generation that enforces distance-based requirements, which for Sic1 creates a heterogeneous compaction and expansion pattern as seen in Supplementary Fig. S6. This property of Sic1, such as overall compactness and large end-to-end distance fluctuations, are consistent with Sic1s ultrasensitive binding to its partner Cdc4. ^28^

**Table 3:** Evaluation of *α*−Syn and Sic1 ensembles from unbiased and experimentally biased IDPForge compared to other methods. Mean absolute errors (MAEs) for J-Couplings (JCs), paramagnetic relaxation enhancement (PREs), chemical shifts (*δ*) and *R_g_* error (Δ*R_g_*) noting experimental uncertainties (*σ_αSyn_* = 3.5*, σ_Sic_*_1_ = 4). Standard deviation (in parenthesis) over 30 ensembles of 100 structures each.

|  | Experimental data type MAE ↓ |  |  |  |  |  |
| --- | --- | --- | --- | --- | --- | --- |
| | JC (Hz) | PRE (Å) | $\Delta R_g$ (Å) | $\delta_{C\alpha}$ (ppm) | $\delta_{C\beta}$ (ppm) | $\delta_H$ (ppm) |
| **α-Syn ensembles** |  |  |  |  |  |  |
| IDPForge Unbiased | 0.347 (0.012) | 4.116 (0.416) | -4.385 (0.331) | 0.231 (0.008) | 0.317 (0.008) | 0.094 (0.002) |
| IDPForge PRE biased | 0.327 (0.011) | **2.565 (0.267)** | -5.941 (0.271) | 0.221 (0.008) | 0.305 (0.008) | 0.094 (0.002) |
| IDPConformerGenerator | 0.292 (0.013) | 4.345 (0.291) | -1.349 (0.665) | 0.623 (0.019) | 0.328 (0.010) | 0.090 (0.003) |
| IDPFold | 0.431 (0.014) | 4.164 (0.289) | -4.304 (0.415) | 0.471 (0.019) | 0.399 (0.009) | 0.103 (0.003) |
| idpGAN | 0.440 (0.010) | 7.864 (0.222) | -9.465 (0.407) | 0.326 (0.005) | 0.473 (0.007) | 0.092 (0.002) |
| STARLING | 0.359 (0.010) | 4.196 (0.314) | 1.960 (0.839) | 0.293 (0.008) | 0.408 (0.009) | 0.086 (0.002) |
| CALVADOS | 0.350 (0.008) | 3.889 (0.207) | 2.582 (1.068) | 0.252 (0.006) | 0.324 (0.006) | 0.093 (0.002) |
| a99SB-disp | 0.351 (0.010) | 5.990 (0.291) | -0.067 (0.870) | 0.228 (0.008) | 0.301 (0.008) | 0.085 (0.003) |
| **Sic1 ensembles** |  |  |  |  |  |  |
| IDPForge Unbiased |  | 8.803 (0.683) | -3.503 (0.407) | 0.588 (0.010) | 0.338 (0.006) | 0.083 (0.003) |
| IDPForge PRE biased |  | **1.978 (0.320)** | -5.498 (0.434) | 0.535 (0.007) | 0.421 (0.009) | 0.086 (0.003) |
| IDPConformerGenerator |  | 6.292 (0.602) | -1.592 (0.580) | 0.759 (0.017) | 0.452 (0.012) | 0.100 (0.004) |
| IDPFold |  | 11.897 (1.281) | 5.149 (0.700) | 0.400 (0.006) | 0.591 (0.007) | 0.103 (0.002) |
| idpGAN |  | 2.037 (0.267) | -3.497 (0.663) | 0.347 (0.008) | 0.741 (0.008) | 0.116 (0.004) |
| STARLING |  | 7.891 (0.669) | 1.265 (0.648) | 0.324 (0.005) | 0.813 (0.007) | 0.119 (0.002) |
| CALVADOS |  | 5.504 (0.724) | -1.856 (0.468) | 0.411 (0.005) | 0.624 (0.008) | 0.102 (0.003) |
| a99SB-disp |  | 2.458 (0.296) | -6.066 (0.531) | 0.598 (0.017) | 0.501 (0.010) | 0.111 (0.007) |

### 2.3 Sampling of IDRs along with their folded domains

Although most disorder in the proteome takes the form of IDRs, at present few approaches are available for modeling IDRs within the context of folded domains at all-atom resolution. For IDPForge, we focus specifically on modeling IDRs by conditioning on available folded structures or structure predictions, rather than attempting to predict folded domains *de novo*. This decision is motivated by several factors: 1) numerous successful structure prediction programs exist for folded proteins, and retraining a model to replicate these results incurs a high computational cost; 2) using existing folded templates allows us to exploit the model architecture to predict disordered regions in a context-aware manner, without requiring additional training; 3) it aligns with the broader objective of unified sequence-based structure prediction for folded and disordered regions.

While IDPForge did not see training examples of IDRs within fixed folded domains, we show the model’s ability to be conditioned on folded template structures by substituting noisy conformations with folded features and applying minimal denoising in those folded domains. We demonstrate IDPForge’s zero-shot generation of terminal and linker IDRs given the structures of folded domains for two cases: modeling the missing electron density from experimentally determined IDR structures (Figure 2), and replacing the low confidence regions longer than 15 amino acids for IDR proteins predicted with AF2 (Figure 3).^1,57^

**Figure 2:**
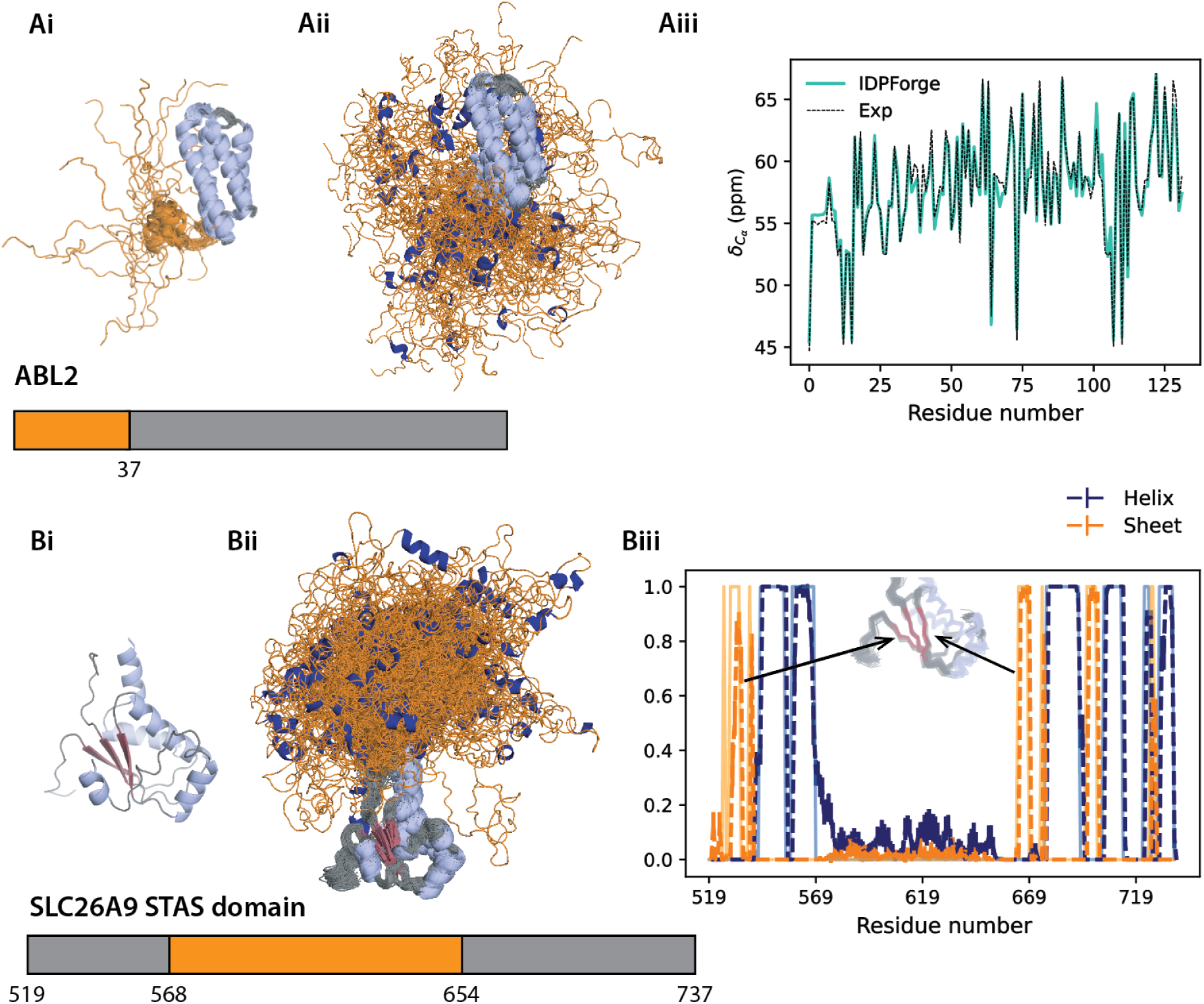
All-atom IDPForge ensembles generated for IDRs with folded domains from experimental structures. Ai) NMR ensemble from PDB ID 2KK1 ^58^. Bi) Intervening 85 residues of the IDR region within the sulfate transporter anti-sigma domain of solute carrier family 26 member 9 (SLC26A9 STAS domain); PDB ID 7CH1 ^59^. (ii) The generated ensembles of ABL2 and SLC26A9 using IDPForge with folded domains shown in grey and IDRs in orange. (Aiii) C*_α_* chemical shifts for IDPForge ABL2 ensemble and solution NMR ^58^. (Biii) Secondary structure propensities for SLC26A9 of the generated ensembles are shown as dark dashed lines and folded templates in light solid colors. Error bars estimate the mean and standard deviation of 100 conformer ensembles from 10 trials. Minor *β*-sheets variations of the modeled folded domains of SLC26A9 STAS domain.

**Figure 3:**
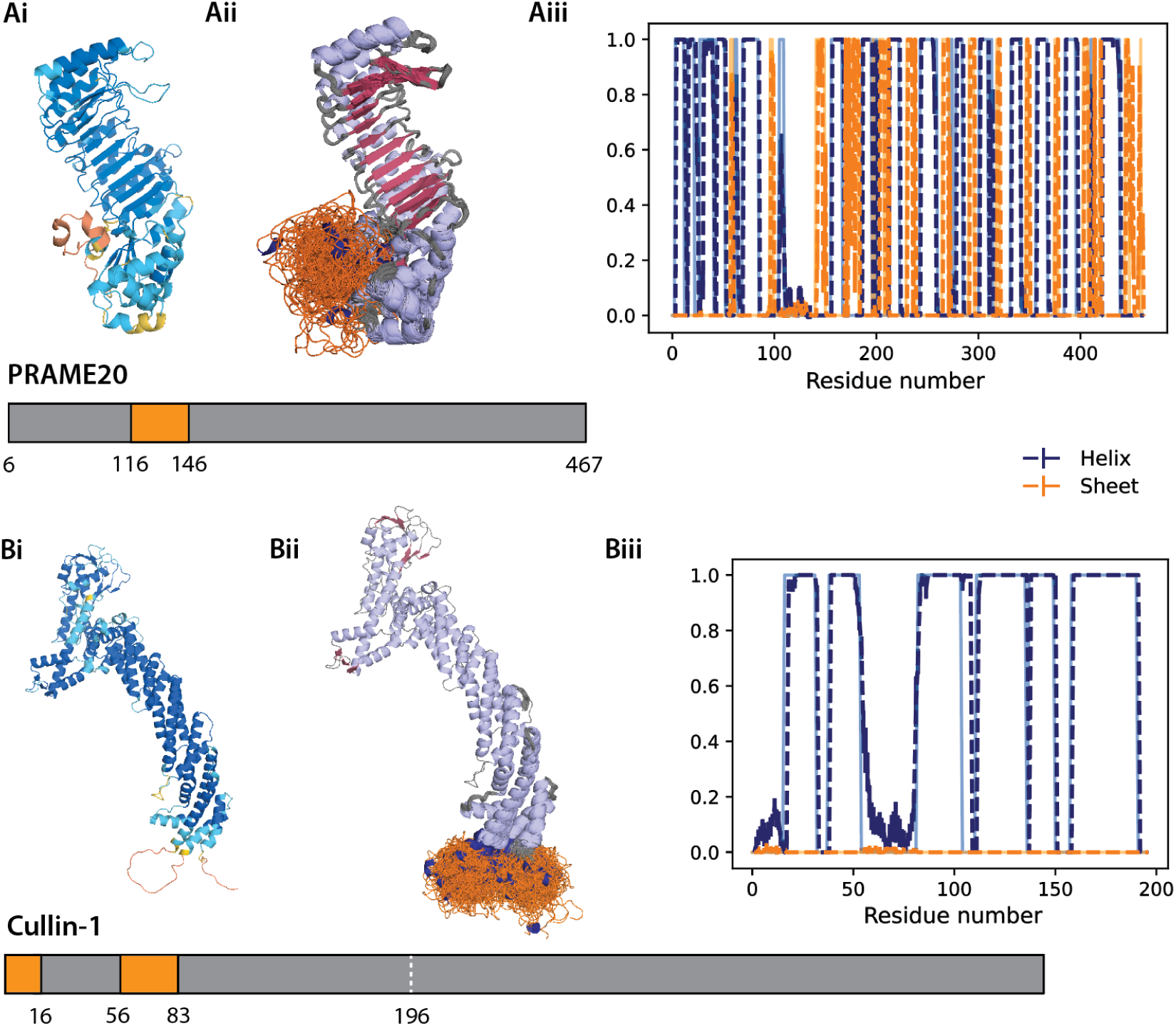
All-atom IDPForge ensembles compared with AlphaFold2 predictions with IDRs representing low confidence regions. A) PRAME family member 20 entry Q5VT98 and B) Cullin-1 entry Q13616 with AF2 predictions (i) colored by pLDDT. Yellow and orange indicate low confidence regions (pLDDT < 70). The generated ensembles of (Aii) PRAME20 and (Bii) Cullin-1 using IDPForge with folded domains shown in grey and IDRs in orange. Secondary structure propensities of (Aiii) PRAME20 and (Biii) Cullin-1 of the generated ensembles (dashed) and the folded templates (solid and in lighter colors).

In the first category, Figure 2Ai considers residues 1-36 in the C-terminal Domain of Tyrosine-protein kinase ABL2 from Homo ^58^ solved with solution NMR, as well as the more difficult case involving 86 unresolved residues in the center of the sulfate transporter anti-sigma (STAS) domain of solute carrier family 26 member 9 (SLC26A9) solved by X-ray crystallography ^59^ shown in Figure 2Bi. We also examine residues 90-123 from human prion protein (HuPrp) with E219K^60^ and Interleukin-6 (IL-6) with 41 missing residues at the N-terminus ^61^ in Supplementary Fig. S7.

Figures 2Aii and 2Bii and corresponding Supplementary Fig. S7ii show the generated ensembles using IDPForge for all four IDR systems, and are seen to be well aligned to the backbone torsion angles from the folded templates while sampling evenly across the alpha, beta and poly-proline II in the disordered regions (shown for IL-6 and STAS in Supplementary Fig. S8). We see that ensemble averages over the IDPForge conformers create good agreement in chemical shifts for the disordered regions while maintaining agreement with experimental CS values in the folded domains (Figures 2Aiii and Supplementary Figs. S7Aiii). The structural metrics show that, like CALVADOS, IDPForge preserves the secondary and tertiary structure in the folded domains, whereas in the disordered regions it is seen that IDPForge predicts more transient secondary structure, unlike CALVADOS which is largely featureless throughout the disordered regions (Figures 2Biii and Supplementary Figs. S7Biii).

In Figure 3Ai we consider the low confidence regions of AF2 structures of the preferentially expressed antigen in melanoma (PRAME) family member 20, replacing the low confidence residues 116-145. We also consider Cullin-1 with two IDR regions: an N-terminal IDR 1-15 and a linker IDR between 56-82 ^62^ in Figure 3Bi. In the latter case we generated the 1-196 residue region for Cullin-1 by first predicting the linker IDR and holding that prediction fixed to then complete the N-terminal IDR generation. Because Cullin-1 beyond residue 196 is distant enough from the IDRs to ignore their influence on sampling, we simply aligned the generated 1-196 regions to the rest of the protein. As expected, IDPForge conformers sample a much greater range of conformations and secondary structure types compared to the primarily featureless coil structures from AF2 for both IDRs (Figures 3ii and 3iii).

We also compare the IDPForge generated ensembles with the multi-domain ensembles modeled by CALVADOS and AFflecto ^47^ across secondary structure categories in Supplementary Fig. S9. AFflecto is a web server that also identifies IDRs from AF predictions and generates ensembles with stochastic sampling and robotic-inspired loop closure. As already seen for the ABL2, HuPrP, IL-6 and STAS IDRs, IDPForge creates more diverse secondary structure types for the disordered regions than those from CALVADOS, while AFflecto exhibits no secondary structure at all.

In and around the sequence junction between the ordered and disordered regions, both IDPForge and CALVADOS exhibit more conformational variations compared to AFflecto as seen in Figure 4. When the junction is helical IDPForge shows more variations across 3_10_-helix, *α_R_*-helix, and *π*-helix states, with some transient helical structure that continues into the beginning of the disordered domain. If the junction involves a folded domain “loop”, as near residue 655 in STAS, more variation is sampled. Evidence of increased conformational sampling near the sequence junctions between folded and disorder regions have been observed previously ^63^, and IDPForge and CALVADOS preserves this feature unlike AFflecto (Figure 4). Both IDPForge and CALVADOS sample the folded domains with average RMSDs to template structures of ≤ 2 Å across all IDRs, whereas AFflecto is seen to introduce very little structural variations in the folded domains with respect to the AF2 predictions (Figure 4). Given the known limitations of the lack of dynamics for AF2 folded structure predictions, we believe that IDPForge and CALVADOS better reflect the true fluctuating ensembles of the folded domains and is a desirable feature of these methods.

**Figure 4:**
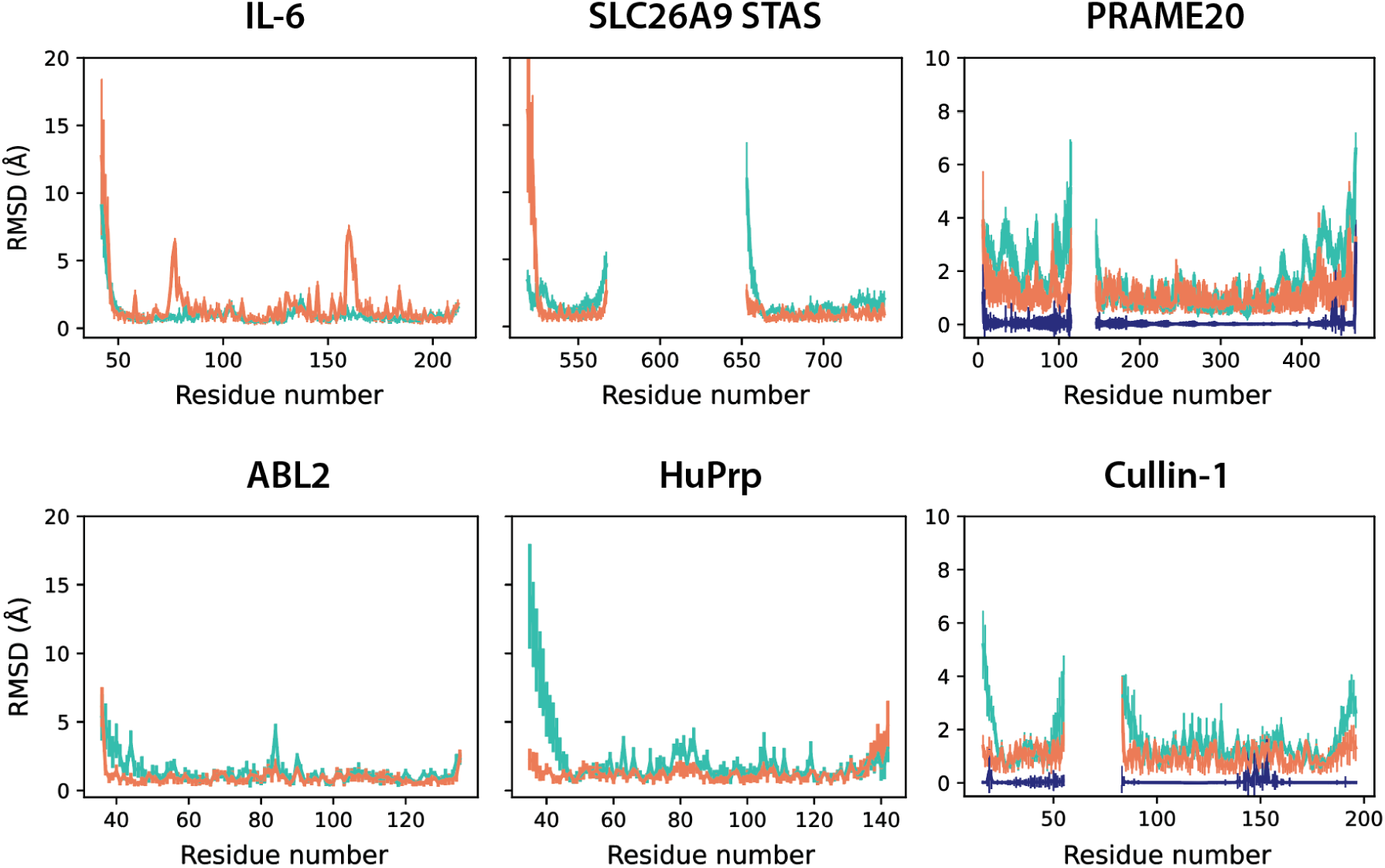
Per residue RMSDs of folded domains for ABL2, HuPrp, IL-6, SLC26A9 STAS, PRAME20 and Cullin-1 ensembles generated by IDPForge, CALVADOS and AFflecto. RMSDs are calculated across ensembles of 100 conformers, based on the folded domain heavy atoms of reference structures from PDB ID 2KK1, 2LFT, 4CNI and 7CH1, and AF2 entry Q5VT98 and Q13616 respectively.

## 3 Discussion and Conclusion

IDPForge is a transformer-based diffusion model that generates both all-atom IDPs and IDRs with folded domains. IDPForge uses attention modules for structure denoising, enabling communication between sequence and pairwise residue information while also providing a suitable channel for conditioning on folded templates. On a benchmark of 32 test IDPs, IDPForge achieves comparable or superior agreement with NMR and SAXS experimental data compared to existing methods. It captures both local structural features as quantified by J-Couplings and global shape characteristics as measured by NOEs, PREs, and the radius of gyration, and benefits from all-atom data training to conform to chemical shift data. We find that other methods show an imbalance among experimental data types from NMR and SAXs, and it would be preferable to have more NMR data to help develop future IDP structural ensemble predictors that perform well on only Rg.

Additionally, its diffusion model framework offers a natural mechanism for incorporating experimental data guidance at inference time. We demonstrate this capability for *α*-Syn and Sic1 in which sampling is guided with PRE data that shifts the conformational distribution towards enhanced agreement for multiple data types with the underlying structural pool better delineating long-ranged residue interactions. We attribute the improvement to the use of the residue frame representation, which enables the model to more effectively integrate distance- and contact-biased experimental restraintsa limitation observed with the torsional representation in DynamICE ^44^. We again highlight that this sampling guidance strategy requires no training but is done at inference, and is generalizable to any sequence with experimental data that can provide direct gradient information.

IDPForge also models regions of disorder within the context of folded domains at inference with a targeted diffusion step strategy by initiating from a structured template instead of a fully noisy state, even though IDPForge is not explicitly trained to perform a partial denoising task. Alternative to stitching based methods that screen a pre-generated pool of single chain conformations ^64^, the attention modules in IDPForge extract and exchange information from both the folded and noisy residue frames through pair and sequence representation for each denoising step. Hence IDPForge can enforce the spatial relation between IDRs and folded domains as well as conditioning on a folded template during generation, an advantage for interpreting linker IDRs. When conditioned on a folded template, IDPForge minimizes perturbations to the structured regions but still introduces local changes in the folded domains and at their junction with IDRs which better reflect an extension of dynamic movements that are present in solution.

There are still limitations to the IDPForge model that will need to be addressed in future work it has not been trained on sequences larger than 200 amino acids and its computational cost scales poorly for larger IDPs and IDRs (Supplementary Figure S10). Even so, there are many exciting directions to use and expand IDPForge over a range of IDP/IDR applications. Because IDPForge does not train a new model for each IDP/IDR sequence, but instead is trained on large variations in sequence data, we anticipate that sequence mutations can be modeled in the disordered domain, just as AF2 is capable of modeling mutations in the folded regions of IDRs. Furthermore,the introduction of PTMs appears to be a general mechanism for regulation of IDPs/IDRs^65^. We can train IDPForge quite easily through data generated not only on canonical amino acids, but phosphorylated sequences given our recent creation of a rotamer library for PTMs ^66^. Finally, given that a significant number of proteins contain IDRs that engage in dynamic interactions with other domains in either discrete dynamic complexes or within condensed states^26,67,68^, future work will pursue simultaneous modeling of multiple IDR domains. In fact, recently IDPForge has been used to create IDR ensembles for ∼200 proteins from the human proteome^69^.

## 4 Methods

### 4.1 Denoising diffusion probabilistic models

Denoising diffusion probabilistic models (DDPMs)^70^ approximate a distribution by parameterizing the reversal of a discrete-time diffusion process. The forward diffusion process increasingly corrupts a sample *x*_0_ from an unknown data distribution *q*(*x*_0_) for a sequence of *T* steps, such that the final step *x_T_* is indistinguishable from a reference distribution that has no dependence on the data. DDPMs approximate the data distribution with a second distribution *p*(*x*_0_) through learning a backward transition kernel *p*(*x_t_*_−1_ | *x_t_*) at each *t*. In the reverse process, one first samples from the reference distribution *x_T_* ∼ *p*(*x_T_*) ≈ *q*(*x_T_*), and then repeatedly denoises by sampling *x_t_*_−1_ ∼ *p*(*x_t_*_−1_ ∼ *x_t_*) until *x*_0_ ∼ *p*(*x*_0_) is obtained.

Under Gaussian noise *ϵ* and *α* that parametrizes a noise schedule (with 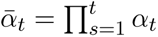), a forward step is defined as,

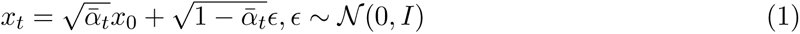

and the conditioned probability in the reverse process as,

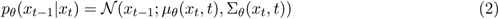

Thus the network can learn the mean of the Gaussian posterior distribution *µ_θ_* from predicting the noise *ɛ_θ_* for the reverse process by

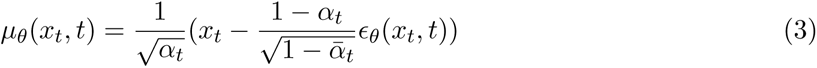

which then allows us to derive a time series of *x* by

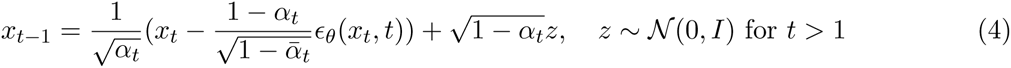

### 4.2 Diffusion over residue rigid frames and torsions

We use DDPMs in the rigid-frame representation of residues, where the forward noising process is defined independently over the rotational and translational components of the backbone rigid bodies, as well as the torsional components of the sidechains. This treatment has been a popular choice for diffusing protein backbones in previous studies^17,71,72^. The diffusion over residue rigid frames is modeled as a discretization of a continuous-time diffusion process with Brownian motion defined on a product manifold T(3) ×*SO*(3) that separates the rotations and translations for each residue^16,21,43^. The translational case is simply defined by a standard Gaussian, whereas the 3D rigid body rotation *SO*(3) can be represented by the isotropic Gaussian distribution ∼*IG_SO_*_(3)_ ^73,74^, which is sampled in the axis-angle parametrization with an angle of rotation *ω* according to

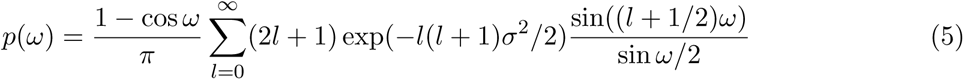

When forward noising the sidechain torsion angles with *SO*(2), we use a wrapped normal distribution where *wrap*(*χ*) = ((*χ* + *π*)*mod*2*π*) − *π*, to handle the periodic nature of torsion angles, a strategy used in FoldingDiff ^19^ and DiffDock ^71^.

While DDPM commonly models the added noise, here we trained the folding block and structure module to predict the final denoised protein structure *x*^_0_. We provide the pseudocode for a training step in Algorithm S1. In this case, a typical denoising step in Eqn. 4 would include a forward network pass and a reverse step that maps the prediction *x*^_0_ to *x_t_*_−1_ given the noised input *x_t_* with independent backward transition kernels on the T(3) × *SO*(3) × *SO*(2) space. The translations *T_t_* and torsions *χ_t_* are updated by interpolating the Gaussian distribution parametrized by the noise schedule *β_t_* = 1 − *α_t_*; this reverse distribution for torsion angles also needs to observe the periodic boundary. On the *SO*(3) space, the backward kernel is approximated with the score matching identity on the corresponding rotations *R_t_* and the predicted noiseless *R*^^^*_t_* such that 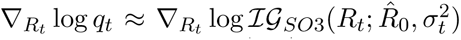. Thus to start a conformer generation, we first randomly sample *x* = (*T, R, χ*^1*,…,*4^)*, T ∼ N*(0*, I*_3_)*, R ∼ SO*(3)*, χ^n^*^=4^ ∼ *wrap*(*N*(0*, I*)) for all disordered residues, and repeatedly apply the denoising step until *T* = 0 (see Algorithm S2). More details of the training scheme and sampling procedures are provided in the Supplementary Text Section 1.1.

### 4.3 Conditioned generation towards experimental data

Conditioned generative models extend the capabilities of generative algorithms by incorporating auxiliary information to meet specific data generation criteria. DDPMs provide a convenient framework for conditioned generation, as the iterative nature of the reverse process offers natural integration points to influence the trajectory of sample synthesis with auxiliary information at each step. In classifier-guided diffusion models, this guidance is achieved by directly incorporating the gradients of a classifier trained to predict the conditional label of interest ∇log *f_ϕ_*(*y x_t_*), to steer the reverse diffusion process toward generating samples that align with condition *y* ^75^. With Bayes’ rule, we can write the log density of the joint distribution *q*(*x_t_, y*) as

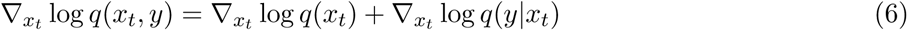

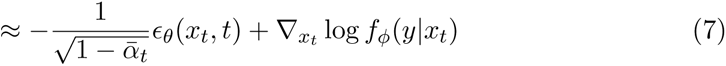

Introducing a guidance scale *ω*, the classifier-guided noise prediction takes the form as,

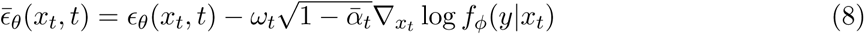

Within the context of generating protein conformers towards desired properties, this is analogous to defining a biasing potential as a function of the structural coordinates *x_t_*_−1_ *x_t_*_−1_ +*ωυ_t_*∇*_x_ P* (*x_t_*). Similar strategies in protein design guided by pseudo-potentials have been adopted by RFdiffusion ^16^ and Chroma ^17^ for tasks such as functional-scaffolding, generating symmetry, or shape constrained protein monomers and oligomers. Here, to bias the generation towards experimental data, we take advantage of the potentials readily defined by the back-calculators. ^38^ Experimental observables such as J-Couplings, NOEs, PREs and *R_g_* (often estimated from a SAXS or smFRET measurement) have direct analytical back-calculations and we limit our studies with experimental guided generation for these data types as no training of an additional classifier is needed. We discuss details of the back-calculators in the Supplementary Text Section 1.1 and 1.3.

### 4.4 Generation of IDRs within the context of folded domains

While IDPForge was originally trained for denoising on completely disordered single chains, we extend IDPForge to infer ensembles of local IDRs with folded domains. This involved conditioning the generation with structural featurizations of the folded domains and their secondary structure assignment in which the model effectively denoises the IDR sequences only by zeroing the diffusion time steps of the folded domains. This partial denoising is controlled by a user defined denoising mask. Inference time modeling of IDRs has the most transferability for generating proteins with a single IDR. Thus, for the case of generating multiple IDRs separated by folded domains, we can model segments of protein containing one IDR at a time and use the denoising mask to treat previous predictions as fixed templates. Sampling of IDRs conditioned on folded domains also requires an estimation of the orientation of the folded domains with respect to the noise initiation. We prepare the folded template input by extracting its coordinates from a reference structure centered by its IDR. We randomly rotate the folded template coordinates and add a noise that scales with the residue number of the IDR to its center of mass. The detailed algorithm is provided in Algorithm S3.

### 4.5 Network architecture

The network architecture illustrated in Figure 1 takes two main module components, the “folding block” and the structure module from ESMFold^45^. The structure module consists of Invariant Point Attention (IPA) followed by backbone update network as introduced in AF2^1^. The structure module is implicitly *SE*(3)-equivariant by frame-localized geometric reasoning. The folding block contains stacks of triangular attention and communications between the sequence and pair representations that closely mimic the Evoformer described in AF2, but with the exception that it does not have a multiple sequence alignment (MSA) processing transformer module. We use two multilayer perceptrons (MLPs): one for the sequence representation to concatenate the diffused sidechain torsion angles and time step encoding, and the other for the pair representation that features a 2D transformation of the diffused coordinates before adding a pairwise relative positional encoding for the residue indices. We then define two learnable embeddings for the protein sequence and for secondary structure encoding (Supplementary Fig. S8), which are added to the sequence MLP outputs before being fed into the attention block. In an ablation study, we also investigated the effect of adding a learnable weighted sum of the ESM2 embedding to the sequence representation or a reshaped ESM2 attention map to the pair representation; we didn’t find significant improvement to IDP ensemble generation performances. Details of these ESM2 appending networks are discussed in the Supplementary Text Section 1.2 and their configurations are listed in the Supplementary Table S2.

### 4.6 Data preparation

We collected the non-overlapping disordered sequences (30-200 residues in length) from the idpGAN training set ^41^ consisting of sequences from DisProt(version 2021-06) ^76^ and the human intrinsically disorder proteome (IDRome) database^33^. We excluded the native folded proteins characterized in the presence of denaturant in the original idpGAN train set that are also in the CASP12 dataset. 32 disordered sequences with experimental data (in Supplementary Table S1) and 6 sequences for systems of folded domains and IDRs (human prion protein (E219K), C-terminal domain of kinase ABL2, interleukin-6, SLC26A9 STAS domain, cullin-1 and PRAME20), were set aside as our test set. In addition to the disordered states, we expanded our data with CASP12 folded states from Sidechainnet^77^ with ≥ 30% thinning, to enrich the training data with secondary structural features. Only CASP12 sequences or segments with completely defined coordinate information and secondary structure annotations were included. We performed sequence similarity clustering with MMseqs2^78^ and removed any sequence with ≥ 30% identity to the test sequences from our trainval set, which in total generated 1,882 disordered sequences and 10,074 CASP12 sequences. We used IDPConformerGenerator ^30^ with sequence-based sampling to generate 100 conformers for each included idpGAN sequences with FASPR^79^ sidechain building. For the IDRome sequences simulated with CALVADOS, we randomly extracted 100 conformers from the coarse-grained trajectories for each sequence, converted to all-atom conformers using cg2all ^80^. All conformers were relaxed for a maximum of 5000 steps with the Amberff14SB forcefield using OpenMM ^81^. We used MDTraj^82^ to assign the secondary structure for each of the generated conformers. This process yielded an extensive structural dataset of 198,399 conformers of which 188,200 corresponded to disordered sequences. For training efficiency, a 90-10% training-validation split was used for CASP12 conformers while a 15-3% split was used for disordered conformers, resulting in a training and validation set comprised of 283 and 57 sequences, respectively.

### 4.7 Timing and memory usage

IDPForge network inference using basic attention calculations on a 132 residue sequence for 40 time steps takes 11.9 seconds per conformer on average on a A40 GPU, in which relaxation takes 8.1 seconds per conformer. Detailed runtime scaling with protein length on a A40 GPU and 32 core CPUs is shown in Figure S9. There can be room for computational efficiency improvements with attention optimization, such as flash attention ^83^ and axial attention chunking, which we did not pursue here.

## 5 Code Availability

The code is available at https://github.com/THGLab/IDPForge.git. X-EISD and UCBShift programs for experimental evaluation are available at https://github.com/THGLab/X-EISD.git and https://github.com/THGLab/CSpred.git. Docker and Apptainer image creation is now outlined in the main README file along with a step-by-step guide for installation on local and hpc cluster environments. We have also added the scoring scripts for reproducibility of Tables 1-3.

## 6 Data Availability

A list of training sequences and all training data and pickle files for secondary structure sampling for IDPForge, as well as all conformer ensembles for each tested method for all 30 test IDPs can be found in Figshare repository https://doi.org/10.6084/m9.figshare.28414937.

## 7 Acknowledgments

All authors acknowledge funding and thank the support from the National Institute of Health under Grant 5R01GM127627-06. J.D.F.-K. also acknowledges support from the Canada Research Chairs Program.

## 8 Author Contributions Statement

O.Z. and T.H.G. designed the project. O.Z. designed the deep leaning network architecture. S.D.C. developed the software and containers for the IDPForge code, and created extensive analysis scripts. Z.-H. L. provided training data using IDPConformerGenerator. O.Z. and T.H.G. wrote the paper. All authors discussed the results and made comments and edits to the manuscript.

## 9 Competing Interests Statement

The authors declare no competing interests.

## Notes

### Competing Interest Statement

The authors have declared no competing interest.

### Summary of Updates

Data Availability. A list of training sequences and all training data and pickle files for secondary structure sampling for IDPForge, as well as all 200 conformer ensembles for each tested method for all 30 test IDPs, can be found in Figshare repository https://doi.org/10.6084/m9.figshare.28414937. Code Availability: The code is available at https://github.com/THGLab/IDPForge.git. X-EISD and UCBShift programs for experimental evaluation are available at https://github.com/THGLab/X-EISD.git and https://github.com/THGLab/CSpred.git. Docker and Apptainer image creation is now outlined in the main README file along with a step-by-step guide for installation on local and hpc cluster environments.

