## Supplementary Info for "IDPForge: Deep Learning of Proteins with Global and Local Regions of Disorder"

### 1 Supplementary Text

#### 1.1 Additional information on diffusion model

**Protein sequence and structural input featurization.** Protein inputs are formatted in either sequence representation or pair representation that is SE(3)-invariant. The sequence representation consists of the amino acid encoding with a padding token, a DSSP secondary structure[1] and Ramachandran combined encoding, a sinusoidal embedding of the time step and the diffused sidechain torsion angles $\chi_t$ in unit vector. The DSSP secondary structure and Ramachandran region combined encoding are defined by $\{H, E, A, B, P, L, C, mask, pad\}$ which correspond to the $\alpha$-helices (H), $\beta$-sheets (E) assigned by DSSP and DSSP-unassigned regions further sub-classed by their Ramachandran regions[2] into right-handed $\alpha_R$ (A), $\beta$ (B), poly-proline II (PPII) (P), left-handed $\alpha_L$ (L) and other coils (C) as shown in Fig. S9. We found that the more detailed Ramachandran encoding helped with all-atom training for the generated structures to adopt more plausible backbone conformations. The *mask* token corresponds to predicting protein conformers with amino sequence dependence and be used as an additional noise injection. The pair representation features diffused backbone coordinates $x_t$ in a stacked map of distogram and anglegram. The distogram is defined by the $C_\beta$ atom (pseudo $C_\beta$ atom interpreted from the $N - C_\alpha - C$ atom positions for glycine) discretized into 33 bins over 2-39 Å with a final bin for distances >39 Å into one-hot encoding. The anglegram is defined by the backbone torsion angles and $C_\beta$ bend and twist. We left-padded all input tensors to the maximum sequence length seen in one batch.

**Algorithm 1** Training

**function** TRAIN(batch)
  **while** Iterate batch **do**
    $t \leftarrow \text{SampleTimestep}(t), t \sim \{1, .., T\}$
    $x_0 \leftarrow batch$
    $x_t = \text{ForwardNoise}(x_0, t)$
    **if** $\text{Uniform}(0, 1) < 0.5$ **or** $t = T$ **then**
      ▷ Train step without self-conditioning
      $\hat{x}_0^{prev} = \vec{0}$
    **else**
      ▷ Train step with self-conditioning
      $x_{t+1} = \text{ForwardNoise}(x_0, t + 1)$
      $\hat{x}_0^{prev} = \text{IDPForge}(x_{t+1}, \vec{0})$
      $\hat{x}_0^{prev} = \text{StopGradient}(\hat{x}_0^{prev})$
    **end if**
    $\hat{x}_0 = \text{IDPForge}(x_t, \hat{x}_0^{prev})$
    Take gradient step on $L(x_0, \hat{x}_0)$
  **end while**
**end function**

**Training.** We present a single training step for the diffusion model in Algorithm 1. We used a linear variance schedule[3] when adding Gaussian noise to the data samples and trained our models with 200 time steps. While one can simply sample time steps uniformly for training, we used a linear scale to put more weights on the larger steps correspond to noisier states such that the model can better handle the more challenging denoising capabilities. Training was split into two phases. The first phase used a 90-10% data split comprised solely of well-ordered CASP12 sequences, while the second phase used a 15-3% split comprised solely of disordered sequences. The preparation of the data was outlined in the Data Preparation section of the main text. Each training phase was conducted on 4 A40 GPUs with bf16 precision, a batch size of 16, and gradient accumulation of 4 for 100 epochs. Additionally, we used an Adam optimizer[4] with a maximum learning rate of 0.001 and gradient clipping of 0.1. The second phase was started immediately after the first phase with the learning rate and epoch gates reset.

**Self-conditioning.** Self-conditioning is a strategy proposed by Chen *et al.*[5] on image generation tasks that recycles the network prediction at the previous time step as an input to subsequent time iterations $\hat{x}_0(x_t, \hat{x}_0^{prev})$ to save calculations for similar successive predictions. When training a model with self-conditioning, we used a 50-50% split for the network taking $\hat{x}_0^{prev}$ as input or no input. When self-conditioning is activated, the network first makes a prediction $\hat{x}_0^{prev}(x_{t+1}, \vec{0})$ without gradient and this prediction is fed back into the network along with $x_t$ to predict $\hat{x}_0(x_t, \hat{x}_0^{prev})$ for loss calculation according to Algorithm 1. The attention block (named as "FoldingTrunk" in ESMFold) refines predictions by recycling sequence and pair outputs with added skip connections of a re-synthesized sequence and pair representation. We input the previous prediction into the network in the same manner only at the start of the attention block, avoiding the need to design additional network architectures.

**Sampling algorithms.** In the denoising process, we define a reverse step that maps $x_{t-1}$ from $\hat{x}_0$ and $x_t$ according to the backward transition kernel in Algorithm 2. The derivation of the kernel on the $SO(3)$ follows[6],

$$\begin{aligned}
\nabla_{R_t} \log q_t &= \mathbb{E}_q[\nabla_{q_t} \log q(x_t|x_0)|x_t] && (1)\\
&\approx \nabla_{R_t} \log q(R_t|R_0 = \hat{R}_0) && (2)\\
&= \nabla_{R_t} \log \mathcal{IG}_{SO3}(R_t; \hat{R}_0, \sigma_t^2) && (3)\\
&= \nabla_{R_t} \omega(\hat{R_t}^\top R_t) \frac{d}{dw} \log p(\omega, \sigma_t^2)|_{\omega=\omega(\hat{R_t}^\top R_t)} && (4)
\end{aligned}$$

The last line denotes a chain rule expansion where $p(\omega, \sigma_t^2)$ refers to the $\mathcal{IG}_{SO(3)}$ sampled in the axis-angle parametrization in Eqn. 4, and $\log \hat{R}^\top R$ is the logarithmic map between $SO(3)$ and the Lie algebra $\mathfrak{so}(3)$, and we refer readers to previous works on diffusion on manifolds[6–8] for more details. Interpolation from this backward transition kernel requires $x_t$ and $\hat{x}_0$ in the same global frame. Thus, we introduce an alignment step to remove the global frame dependence. When a folded domain (not denoised) is present, the alignment can be simply done between $\hat{x}_0$ and $x_t$ using the fixed domain as reference (in Algorithm 3); otherwise we align $\hat{x}_0$ to $x_t$. This allows the use of a SE(3) invariant loss as discussed in the loss function section below.

To condition generation on Ramachandran regions and/or DSSP propensities, a sampling of the tokens can be done based on the user-defined data or adhering to ones inferred from chemical shift data using softwares such as $\delta$2D[9] and CheSPI[10]. We report ensembles with random sampling of the backbone Ramachandran probability distributions of short peptides by sliding over

**Algorithm 2** Sampling

**function** ReverseStep($x_t, \hat{x}_0$)
  $(R_t, T_t, \chi_t) \leftarrow x_t$
  $(\hat{R}_0, \hat{T}_0, \hat{\chi}_0) \leftarrow \hat{x}_0$
  Update translations:
  $T_{t-1} \sim \mathcal{N}(\sqrt{\bar{\alpha}_{t-1}} \frac{1-\alpha_t}{1-\bar{\alpha}_t} \hat{T}_0 + \sqrt{\alpha_t} \frac{1-\bar{\alpha}_{t-1}}{1-\bar{\alpha}_t} T_t, \beta_t)$
  Update rotations:
  $\boldsymbol{s} \leftarrow \text{RotationScoreApproximation}\left(R_t, \hat{R}_0, \sigma_t^2\right)$
  $\Delta R \leftarrow \exp\left\{(\sigma_t^2 - \sigma_{t-1}^2)(R_t)^\top \boldsymbol{s}\right\}$ ▷ Exponential map from Lie algebra to $SO(3)$
  $R_{t-1} \sim \mathcal{IG}_{SO(3)}\left(R_t \Delta R, \sigma_t^2 - \sigma_{t-1}^2\right)$
  Update torsions:
  $\chi_{t-1} \sim \mathcal{N}_{wrap}(\sqrt{\bar{\alpha}_{t-1}} \frac{1-\alpha_t}{1-\bar{\alpha}_t} \hat{\chi}_0 + \sqrt{\alpha_t \frac{1-\bar{\alpha}_{t-1}}{1-\bar{\alpha}_t}} \chi_t, \beta_t)$ ▷ Wrap about $[-\pi, \pi)$
  $x_{t-1} \leftarrow (R_{t-1}, T_{t-1}, \chi_{t-1})$
  **return** $x_{t-1}$
**end function**

**function** Sample($L$)
  ▷ Generation of $L$-residue structure
  $x_T \leftarrow \text{RandomSample}(L)$
  $\hat{x}_0^{prev} = \vec{0}$
  **for** $t = T, \ldots, 1$ **do**
    $\hat{x}_0 = \text{IDPForge}(x_{t+1}, \hat{x}_0^{prev}$ if self-conditioning else $\vec{0})$
    $\hat{x}_0 \leftarrow \text{Align}(\hat{x}_0, x_t)$
    $x_{t-1} \leftarrow \text{ReverseStep}(x_t, \hat{x}_0)$
    $\hat{x}_0^{prev} = \hat{x}_0$
  **end for**
  **return** $\hat{x}_0$
**end function**

and matching consecutive sequence fragments extracted from the training data. This strategy is inspired from IDPConformerGenerator[11] and the Ramachandran sampling database can also be customized by curating from desired PDB source. For the folded domains, we calculate its DSSP assignments as the conditioning input.

While typically the same sequence of diffusion time steps is used in training and inference, reducing the number of steps in inference is desirable to improve sampling speed and thus the applicability in generating larger protein structures. To reduce the number of sampling steps from $T = 200$ to $T' = 40$, we scale the variance schedule such that the noise added in each new time step $t'$ are proportionally to the implied fraction of the trajectory traversed over the training $T$ steps.

**Algorithm 3** Generation with experimental guidance and local IDRs with folded domains

**function** SampleGuided($L, P, \omega$)
  ▷ Generation of $L$-residue structure, guided by potential $P$
  $x_T \leftarrow$ RandomSample($L$)
  **for** $t = T, \ldots, 1$ **do**
    $\hat{x}_0 \leftarrow$ IDPForge($x_t$)
    $\hat{x}_0 \leftarrow \hat{x}_0 + \omega_t \nabla_{x_0} P(\hat{x}_0))$ ▷ Apply guidance
    $x_{t-1} \leftarrow$ ReverseStep($x_t, \hat{x}_0$)
  **end for**
  **return** $\hat{x}_0$
**end function**

**function** SampleLDRs($L, x^{folded}$)
  ▷ Generation of $L$-residue structure including local IDRs and template folded structures
  $x_T \leftarrow$ RandomSample($L^{IDR}$)
  $t^{l \in \{folded\}} = 0$
  $x_T^{l \in \{folded\}} = x^{folded}$
  **for** $t^{l \in \{IDR\}} = T, \ldots, 1$ **do**
    $\hat{x}_0 \leftarrow$ IDPForge($x_t$)
    $\hat{x}_0 \leftarrow$ Align($x_t, \hat{x}_0 | x^{folded}$) ▷ Align based on the folded domains
    $x_{t-1} \leftarrow$ ReverseStep($x_t, \hat{x}_0 | x^{l \in \{IDR\}}$) ▷ Only apply to IDRs
  **end for**
  **return** $\hat{x}_0$
**end function**

A sampling step guided with a user-defined potential for the diffusion model is shown in Algorithm 3. Here, the potentials can be simply formulated based on the back-calculations of the experimental types of interest, which can be found in the X-EISD work[12]. Given that these back-calculators are defined as functions of completely denoised protein structures and may not be valid for intermediate noisy coordinates, we apply the guidance step at the prediction of $\hat{x}_0$ and before interpolation to $x_{t-1}$. In general, we use a L2 loss between the experimental data and the ensemble back-calculated observables as the biasing potentials. To account for the generous experimental uncertainties of NOEs and PREs when interpreted as distance restraints, the distance restraints take the form of a flat-bottomed potential,

$$P(\hat{x}_0, d_{exp}, \xi) = \begin{cases} (d(\hat{x}_0) - d_{exp} + \xi)^2, & \text{if } d(\hat{x}_0) < d_{exp} - \xi, \\ 0, & \text{if } d_{exp} - \xi \leq d(\hat{x}_0) \leq d_{exp} + \xi, \\ (d(\hat{x}_0) - d_{exp} - \xi)^2, & \text{if } d(\hat{x}_0) > d_{exp} + \xi. \end{cases} \tag{5}$$

**Loss function.** Frame Aligned Point Error (FAPE) loss was introduced in AF2[13] and is broadly used by structure prediction models.[14–16] It exhibits good performance as its SE(3) invariance keeps the chirality of relevant atomic centers in the protein. While it has been previously applied to a diffusion model[17], other works have indicated that its direct use may be inappropriate under a matching objective[6,18]. To reconcile FAPE loss with the diffusion framework, we use the Fréchet mean of FAPE as proposed by Jing *et al.*[18],

$$L = \text{FAPE}^2(\hat{x_0}, x_0) \tag{6}$$

where the protein structures are redefined in the quotient space $\mathbb{R}^{3\times N}/SE(3)$ that eliminates global frame dependence after alignment. We defined auxiliary loss functions in addition to the squared FAPE loss,

$$L = L_{\text{FAPE}^2} + c_{ang}L_{ang} + c_{dist}L_{dist} + c_{viol}L_{viol} \tag{7}$$

which include angular loss to minimize the distance before the predicted and target torsion angles while enforcing the predictions as unit vectors,

$$L_{ang} = ||\vec{\theta} - \hat{\vec{\theta}}|| + 0.02L_1(||\hat{\vec{\theta}}|| - 1) \tag{8}$$

an inter-residue distance loss defined based on the $C_\beta$ atom,

$$L_{dist} = MSE(d_{C_\beta} - \hat{d}_{C_\beta})_{clamp} \tag{9}$$

where we clamped the distance error at 10Å; for residues assigned as helices or sheets, we define a one-sided clamp to encourage learning the contacts. We applied a structural violation loss to penalize atomic distances that violate covalent bond definitions and van der Waal radii, only at later stages of training as the structural violation term is computationally expensive and can cause training instability[13].

#### 1.2 Ablation study on ESM evolutionary embedding

ESM2 (Evolutionary Scale Modeling 2) provides rich embeddings that capture the evolutionary, structural, and functional context of proteins, and can inform protein modeling without the need to perform a multiple sequence alignment. We explored two forms of ESM to include into our network: 1) the embedding output from ESM2, which is summed with learnable weights and then concatenated along with the torsion angles and time step encoding before passed to the sequence processing MLP network; 2) a flattened ESM2 attention map that is first condensed by a MLP before added to the pair representation of the diffused residue frames. We denote the 2 models as *model ESM seq* and *model ESM pair*, as compared with original model, *model no ESM embed*, in the following analysis. The *model no ESM embed* only provides additional input but no trainable weights. Using model version esm2_8M_270K, we trained all models in the same manner with hyperparameters listed in Table S2. As no significant model improvement is observed for the models augmented with ESM embeddings but with more parameters, we focused on *model no ESM embed* refered to as IDPForge for additional IDPs/IDRs generation tasks.

#### 1.3 Ensemble preparation and characterization

**Preparation.** We compare IDPForge performance with 6 other methods, representative of the 3 main approaches for generating disordered protein ensembles: all-atom and coarse-grained force-fields, statistical/knowledge-based methods, and ML models. While there are other published methods for relevant tasks, our selection is limited by accessibility of their software implementations. We reported idpGAN[19] instead of idpSAM[20] as the open source weights of idpSAM predict overly compact ensembles, which is also noted in the supplementary analysis of Ref[21] and[22]. We extracted 500 conformers evenly across 9 disordered proteins from all-atom trajectories with a99SB-disp from Ref[23] and kindly released to us. For CALVADOS simulations[24], we sampled trajectories for all 30 test sequences and the 6 IDRs with folded domains systems with 295.15 K, pH=6.5 and an ionic strength derived from 150mM NaCl solution. Among the methods we compare with, CALVADOS, idpGAN, STARLING and IDPFold generates coarse-grained or backbone-only ensembles, which we mapped to all-atom representations for compatibility. We used cg2all[25] for CG-to-AA conversion and pyRosetta[26] to populate the sidechain states for IDPFold, both followed by energy minimization with the Amberff14SB forcefield[27] using OpenMM[28]. We note that minimization is necessary for evaluation as the direct outputs from all-atom mappings sometimes have local structural violations that can induce error for back-calculations, as was also the case in AF2[13]. We also consider CALVADOS and AFflecto[29] for comparison of performances to generate IDRs and folded domains ensembles. While AFflecto assigns IDR boundaries based on pLDDT scores and residue contact numbers, we manually set the IDR boundaries to include low confidence regions longer than 15 amino acids. For consistency, we find that subsampling from a pool of atleast 200 conformers leads to standard deviations within experimental noise, so 200 conformers are generated for each method per protein.

**X-EISD calculation.** The X-EISD method applies a maximum likelihood estimator to formulate a log likelihood as the degree to which a simulated ensemble is in agreement with a set of experimental data, given both the experimental and back-calculation uncertainties modeled as optimized Gaussian random variables under a Bayesian framework. X-EISD can be applied to generated an aggregated score of multiple data types as shown in Eq.10,

$$logp(X, \xi|D, I) = logp(X|I) + \sum_{j=1}^{M} log[p(d_j|X, \xi_j, I)p(\xi_j|I)] + C \tag{10}$$

where X is a set of conformers, $\xi$ denotes the various uncertainties, D is the experimental data and I is any other prior information. We refer readers to references[12,30] for more detailed descriptions of the approach. The preparations of experimental data and back calculation uncertainties for the reported data types are also described previously[12], but summarized here: the experimental (exp) and back calculations (bc) errors for chemical shifts $\sigma^{exp}_{C\alpha,C\beta}$=0.3 ppm, $\sigma^{exp}_{H}$=0.03 ppm; $\sigma^{bc}_{C\alpha}$=0.97 ppm, $\sigma^{bc}_{C\beta}$=1.26 ppm, $\sigma^{bc}_{H}$=0.38 ppm; JCs $\sigma^{exp}$=0.5 Hz; $\sigma^{bc}_{A}$=0.14 Hz, $\sigma^{bc}_{B}$=0.03 Hz, $\sigma^{bc}_{C}$=0.08 Hz; PREs $\sigma^{exp}$=5.0 Å; $\sigma^{bc}$=0.001 Å; smFRET$\langle E\rangle$ $\sigma_{exp}$=0.02; $\sigma_{bc}$=0.0074. We use UCBShift[31] for chemical shift calculations, and other experimental data types such as J-couplings, NOEs, PREs and the efficiencies of smFRETs are treated using in-house scripts. We normalized the aggregated X-EISD score based on the range of the scores for each protein by different models,

$$X_{norm} = (X - X_{min})/X_{max} \tag{11}$$

prior to reporting an averaged ranking score for based on all proteins.

**Evaluation metrics.** In addition to X-EISD scores, we also report chemical shift $\chi^2$, taking into account of the uncertainties of the back-calculator errors for different atom types,

$$\chi^2 = \frac{1}{N}\sum_i \frac{(\delta_i^{exp} - \delta_i)^2}{\sigma_i^2} \tag{12}$$

where $\chi^2 \leq 1$ can be viewed as achieving a reasonable fit, though the assumptions that the weighted residuals are standard normally distributed required for $\chi^2$ statistics are only approximately held. We also reported mean absolute error (MAE) to quantify the agreement between back-calculations and experimental data. For NOEs, PREs and $R_g$ errors ($\Delta R_g$), we used the same treatment as in Eqn. 5 to exclude errors within uncertainty ranges.

**Structural validation.** We validate IDPForge to generate physical protein conformers by examining the torsion angle distributions and the inter-residue bond angles and bond lengths. In Fig. S1, we show that the backbone torsion distribution follows a typical Ramachandran plot and sidechain angle densities are also consistent with the training data. Since the model diffuses the protein structure by each residue frame, we confirm that IDPForge can capture the correct connectivity encoded by C-N bond lengths, CA-C-N angles and $\omega$ torsion angles, although the slightly broad distribution can be easily resolved with an energy minimization step. We also relax the generated structures with Amberff14SB forcefield[27] for a maximum of 20 L-BFGS minimization attempts to resolve sidechain collisions.

#### 2 Supporting Tables

**Table S1: Sequences of test set IDPs with experimental data.** Data types include radius of gyration $R_g$, $^3J_{H_N H\alpha}$ (JC), backbone chemical shifts (CS), NOE, PRE and smFRET. When multiple experimental $R_g$ were reported in literature, we take the mean and error range from these values. * marked sequences are in the CALVADOS, IDPFold, and/or idpGAN training sets.

| Name | Sequence | Length | $R_g$(Å) | Other Exp. data | Ref. |
|---|---|---|---|---|---|
| RS1* | GAMGPSYGRSRSRSRSRSRSRSRS | 24 | 12.6±0.1 | JC, CS | [32] |
| His5* | DSHAKRHHGYKRKFHEKHHSHRGY | 24 | 13.8±0.3 | JC, CS | [33] |
| N49 | GCQTSRGLFGNNNTNNINNSSSGMNNASAGLFGSKP | 36 | 16.3±1.0 | | [34] |
| A$\beta$40* | DAEFRHDSGYEVHHQKLVFFAEDVGSNKGAIIGLMVGGVV | 40 | 13.3±2.3 | JC, NOE, CS | [35–38] |
| NLS | ACETNKRKREQISTDNEAKMQIQEEKSPKKKRKKRSSKANKPPE | 44 | 24.0±3.0 | | [34] |
| drkN-SH3 | MEAIAKHDFSATADDELSFRKTQILKILNMEDDSNWYRAELDGKEGLIPSNYIEMKNHD | 59 | 18.3±1.6 | JC, NOE, PRE, smFRET, CS | [39–42] |
| N-WASP | GAMGTAGNKAALLDQIREGAQLKKVEQNSRPVSCSGRDALLDQIRQGIQLKSVADGQESTPPTPAPT | 67 | 24.3±0.43 | | [43] |
| ACTR** | GTQNRPLLRNSLDDLVGPPSNLEGQSDERALLDQLHTLLSNTDATGLEEIDRALGIPELVNQGQALEPKQD | 71 | 25.0±1.0 | CS | [28,44,45] |
| PaaA2* | MDYKDDDDKNRALSPMVSEFETIEQENSYNEWLRAKVATSLADPRPAIPHDEVERRMAERFAKMRKERSKQ | 71 | 22.4±4.0 | CS | [46] |
| IB5* | SARSPPGKPQGPPQQEGNKPQGPPPPGKPQGPPPAGGNPQQPQAPPAGKPQGPPPPPQGGRPPRPAQGQQPPQ | 73 | 32.0±2.0 | | [47] |
| PIR domain* | SVSPMRSVSENSLVAMDFSGQKTRVIDNPTEALSVAVEEGLAWRKKGCLRLGNHGSPTAPSQSSAVNMALHRSQP | 75 | 26.5±1.0 | | [48] |
| NUS** | PSASPAFGANQTPTFGQSQGASQPNPPGFGSISSSTALFPTGSQPAPPTFGTVSSSSQPPVFGQQPSQSAFGSGTTPN | 78 | 25.0±1.0 | | [34] |
| Ash1** | GASASSSPSPSTPTKSGKMRSRSSSPVRPKAYTPSPRSPNYHRFALDSPPQSPRRSSNSSITKKGSRRSSGSSPTRHTTRVCV | 83 | 28.5±3.4 | CS | [49] |
| Sic1* | GSMTPSTPPRSRGTRYLAQPSGNTSSSALMQGQKTPQKPSQNLVPVTPSTTKSFKNAPLLAPPNSNMGMTSPFNGLTSPQRSPFPKSSVKRT | 92 | 30.0±4.0 | PRE, CS | [50,51] |

Table S1 (continued)

| Name | Sequence | Length | $R_g$(Å) | Other Exp. data | Ref. |
|---|---|---|---|---|---|
| p53 1-93* | MEEPQSDPSVEPPLSQETFSDLWKLLPENNVLSPLPSQAMDDLMLSPDDIEQWFTEDPGPDEAPRMPEAAPPVAPAPAAPTPAAPAPAPSWPL | 93 | 28.7±1.0 | CS | 52,53 |
| IBB | GCTNENANTPAARLHRFKNKGKDSTEMRRRRIEVNVELRKAKKDDQMLKRRNVSSFPDDATSPLQENRNNQGTVNWSVDDIVKGINSSNVENQLQAT | 97 | 32.0±2.0 | | 34 |
| Tau K19** | QTAPVPMPDLKNVKSKIGSTENLKHQPGGGKVQIVYKPVDLSKVTSKCGSLGNIHHKPGGGQVEVKSEKLDFKDRVQSKIGSLDNITHVPGGGNKKIE | 98 | 25.0±2.0 | | 54 |
| HIV-1Tat$_{133}$* | MEPVDPRLEPWKHPGSQPRTACTNCYCKKCCFHCQVCFIRKALGISYGRKKRRQRRRAPPDSETHQVSPPKQPASQPRGDPTGPKESKKKVERETETHPVN | 101 | 33.0±1.5 | | 55 |
| p15$^{PAF}$** | VRTKADSVPGTYRKVVAARAPRKVLGSSTSATNSTSVSSRKAENKYAGGNPVCVRPTPKWQKGIGEFFRLSPKDSEKENQIPEEAGSSGLGKAKRKACPLQPDHTNDEKE | 110 | 28.1±0.3 | CS | 56 |
| ProT$\alpha$ | MSDAAVDTSSEITTKDLKEKKEVVEEAENGRDAPANGNAENEENGEQEADNEVDEEEEEGGEEEEEEEEGDGEEEDGDEDEEAESATGKRAAEDDEDDDVDTKKQKTDEDD | 111 | 37.8±0.9 | | 57 |
| NUL* | GCGFKGFDTSSSSSNSAASSSFKFGVSSSSSGPSQTLTSTGNFKFGDQGGFKIGVSSDSGSINPMSEGFKFSKPIGDFKFGVSSESKPEEVKKDSKNDNFKFGLSSGLSNPV | 112 | 30.0±3.0 | | 34 |
| KISS-1$^{19-138}$* | GEPLEKVASVGNSRPTGQQLESLGLLAPGEQSLPCTERKPAATARLSRRGTSLSPPPESSGSPQQPGLSAPHSRQIPAPQGAVLVQREKDLPNYNWNSFGLRFGKREAAPGNHGRSAGRG | 120 | 34.7±0.5 | CS | 58 |
| ERM* | MDGFYDQQVPFMVPGKSRSEECRGRPVIDRKRKFLDTDLAHDSEELFQDLSQLQEAWLAEAQVPDDEQFVPDFQSDNLVLHAPPPTKIKRELHSPSSELSSCSHEQALGANYGEKCLYNYCA | 122 | 39.6±0.7 | | 59 |
| RNaseA* | KETAAAKFERQHMDSSTSAASSSNYCNQMMKSRNLTKDRCKPVNTFVHESLADVQAVCSQKNVACKNGQTNCYQSYSTMSITDCRETGSSKYPNCAYKTTQANKHIIVACEGNPYVPVHFDASV | 124 | 33.6±0.1 | | 60 |
| $\gamma$-Synuclein | MDVFKKGFSIAKEGVVGAVEKTKQGVTEAAEKTKEGVMYVGAKTKENVVQSVTSVAEKTKEQANAVSEAVVSSVNTVATKTVEEAENIAVTSGVVRKEDLRPSAPQQEGEASKEKEEVAEEAQSGGD | 127 | | CS | 61 |
| Tau K18** | QTAPVPMPDLKNVKSKIGSTENLKHQPGGGKVQIINKKLDLSNVQSKCGSKDNIKHVPGGGSVQIVYKPVDLSKVTSKCGSLGNIHHKPGGGQVEVKSEKLDFKDRVQSKIGSLDNITHVPGGGNKKIE | 129 | 38.0±3.0 | | 54 |
| N$_{TAIL}$* | MHHHHHHTTEDKISRAVGPRQAQVSFLHGDQSENELPRLGGKEDRRVKQSRGEARESYRETGPSRASDARAAHLPTGTPLDIDTASESSQDPQDSRRSADALLRLQAMAGISEEQGSDTDTPIVYNDRNLLD | 132 | 27.5±0.4 | CS | 62,63 |

Table S1 (continued)

| Name | Sequence | Length | $R_g$(Å) | Other Exp. data | Ref. |
|---|---|---|---|---|---|
| $\beta$-Synuclein | MDVFMKGLSMAKEGVVAAAEKTKQGVTEAAEKTKEGVLYVGSKTREGVVQGVASVAEKTKEQAS HLGGAVFSGAGNIAAATGLVKREEFPTDLKPEEVAQEAAEEPLIEPLMEPEGESYEDPPQEEYQEY EPEA | 134 | | CS | [64] |
| $\alpha$-Synuclein* | MDVFMKGLSKAKEGVVAAAEKTKQGVAEAAGKTKEGVLYVGSKTKEGVVHGVATVAEKTKEQVT NVGGAVVTGVTAVAQKTVEGAGSIAAATGFVKKDQLGKNEEGAPQEGILEDMPVDPDNEAYEMPSE EGYQDYEPEA | 140 | 35.5±3.5 | JC, PRE, smFRET, CS | [65–72] |
| FhuA* | ESAWGPAATIAARQSATGTKTDTPIQKVPQSISVVTAEEMALHQPKSVKEALSYTPGVSVGTRG ASNTYDHLIIRGFAAEGQSQNNYLNGLKLQGNFYNDAVIDPYMLERAEIMRGPVSVLYGKSSPG GLLNMVSKRPTTEPL | 143 | 33.4±0.2 | | [60] |
| N98** | GCFNKSFGTPFGGGTGGFGTTSTFGQNTGFGTTSGGAFGTSAFGSSNNTGGLFGNSQTKPGGLF GTSSFSQPATSTSTGFGFGTSTGTANTLFGTASTGTSLFSSQNNAFAQNKPTGFGNFGTSTSSGGL FGTTNTTSNPFGSTSGSLFGP | 151 | 29.5±1.0 | | [34] |
| NSP | GCNFNTPQQNKTPFSFGTANNNSNTTNQNSSTGAGAFGTGQSTFGFNNSAPNNTNNANSSITPA FGSNNTGNTAFGNSNPTSNVFGSNNSTTNTFGSNSAGTSLFGSSSAQQTKSNGTAGGNTFGSSSLF NNSTNSNTTKPAFGGLNFGGGNNTTPSSTGNANTSNNLFGATANAN | 176 | 40.5±0.5 | | [34] |

**Table S2: Network configuration and hyperparameters.**

| | IDPForge (model no ESM embed) | model ESM seq | model ESM pair |
|---|---|---|---|
| $C_t$ | 32 | 32 | 32 |
| ESM seq combine dim | - | 7 | - |
| ESM pair MLP input dim | - | - | 120 |
| $C_s^{MLP}$ | 40 | 360 | 40 |
| $C_z^{MLP}$ | 71 | 71 | 71 |
| Folding block | | | |
| $C_s$ | | 128 | |
| $C_z$ | | 64 | |
| attention head width | | 32 | |
| max recycles | | 3 | |
| n blocks | | 2 | |
| Structural module | | | |
| $C_s$ | | 256 | |
| $C_z$ | | 64 | |
| n blocks | | 4 | |
| Loss | | | |
| FAPE clamp distance | | 10 | |
| FAPE clamp ratio | | 0.5 | |
| $c_{ang}$ | | 0.1 | |
| $c_{dist}$ | | $0.01^{epoch>50}$ | |
| $c_{viol}$ | | $0.01^{epoch>80}$ | |

### 3 Supporting Figures

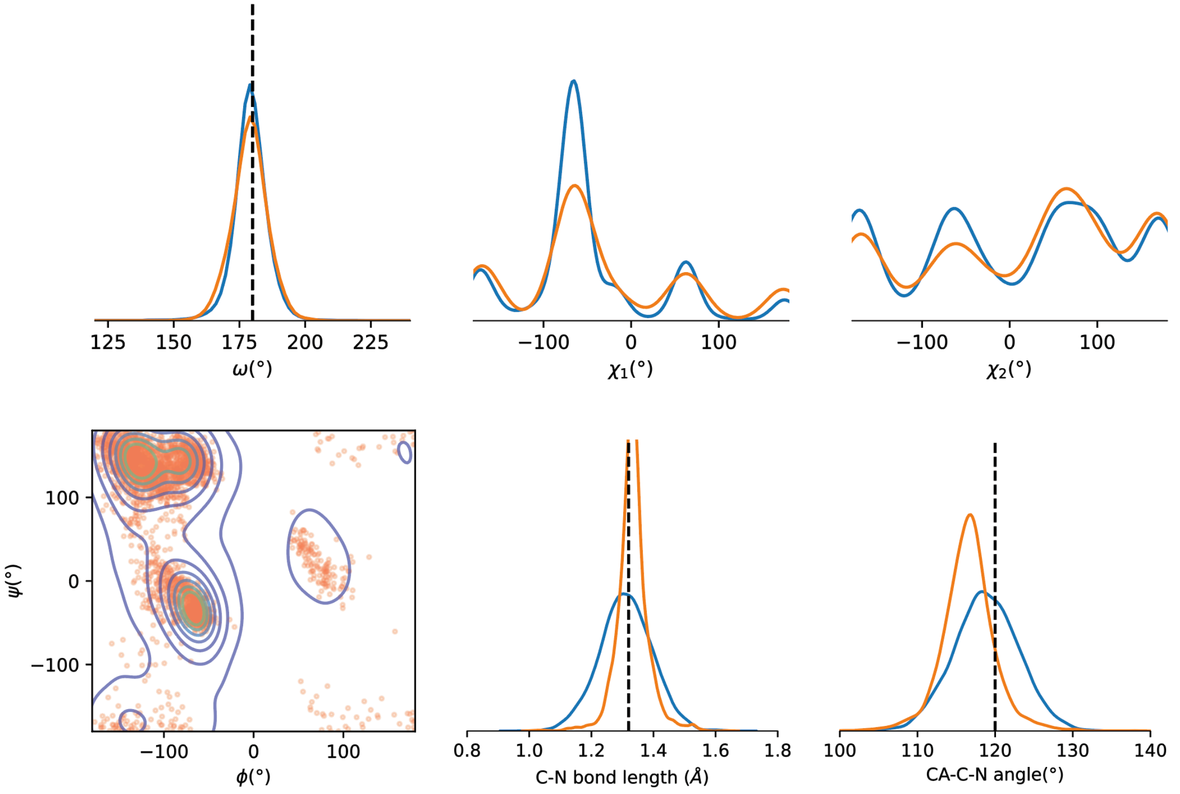


**Figure S1: Bond, angle and torsion statistics of IDPForge sampled conformations.** All plots are color coded to denote the source of the data (Blue: IDPForge, Green: CASP12 Pre-training data, Orange: IDP Fine-tuning data). Additionally, IDPForge is shown in contour and the training data in scattering for the torsion plot.

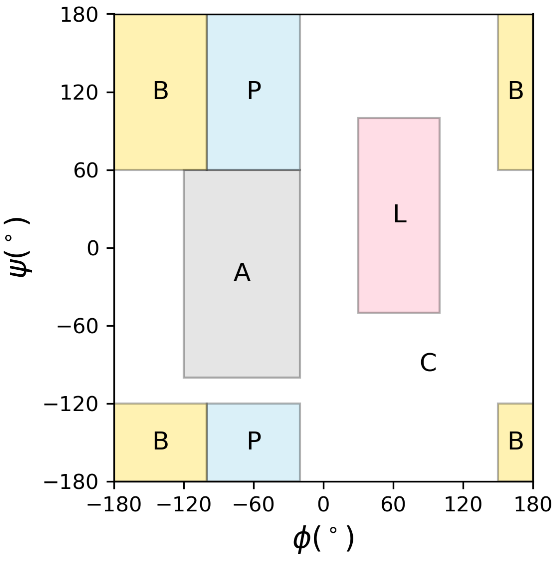


**Figure S2: Definition of Ramachandran region encoding.**

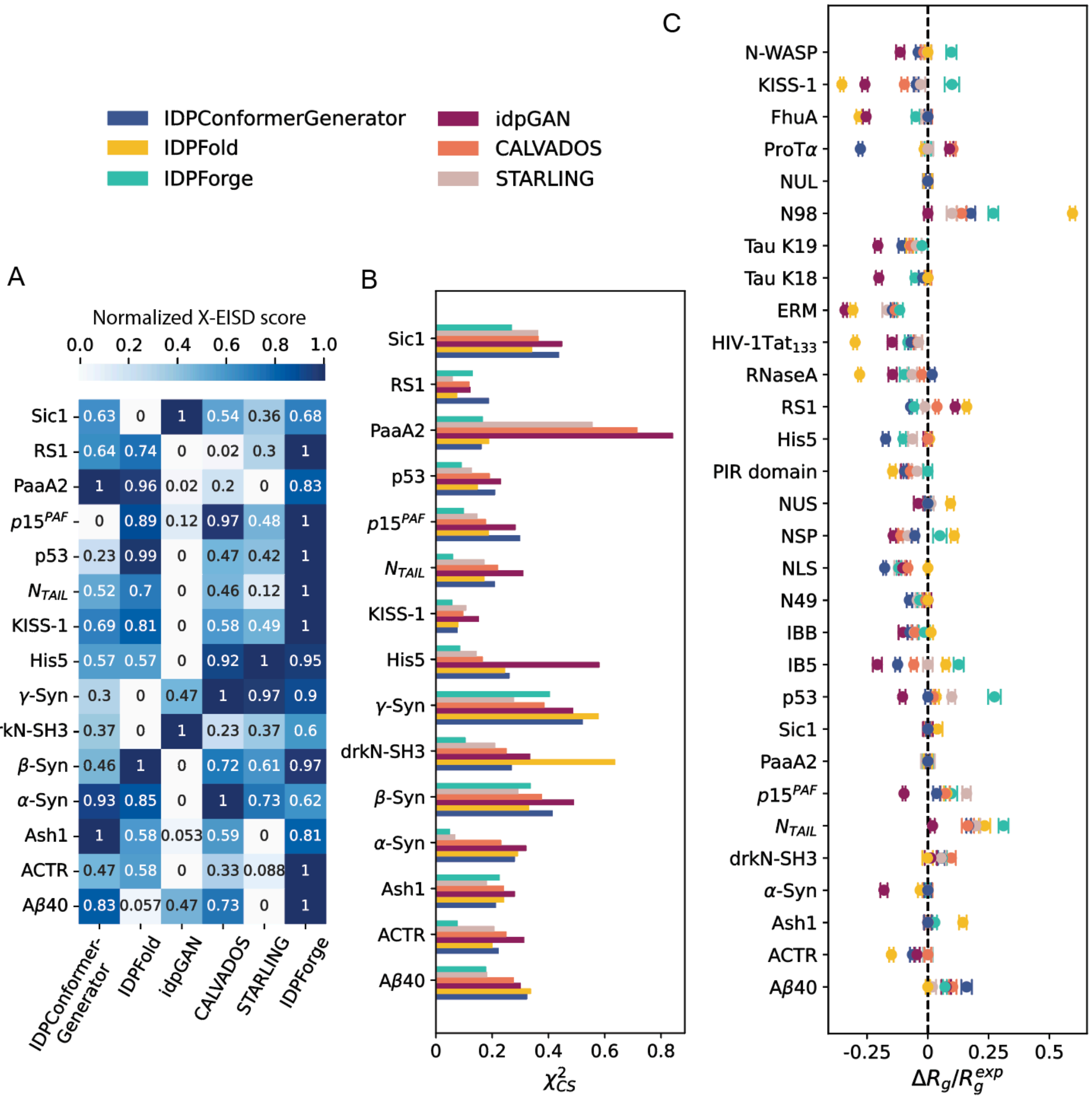


**Figure S3: Evaluation of the generated ensembles for the test sequences for various methods.** (A) Aggregate X-EISD score over all available data types normalized per protein; X-EISD scores range between 0-1, and the higher score indicates better experimental agreement. (B) Chemical shift $\chi^2$ for all assigned atom types with consideration of the back-calculation errors (Eqn. S10) for 15 IDPs. (C) Normalized radius gyration error with respect to the experimental values for 30 IDPs. Experimental uncertainties were accounted for by $\Delta R_g = max(\langle R_g \rangle - R_g^{exp} - \sigma^{exp}, 0) + min(\langle R_g \rangle - R_g^{exp} + \sigma^{exp}, 0)$ (also see Eqn. S5). Compaction: $\Delta R_g / R_g^{exp} < 0$ and expansion: $\Delta R_g / R_g^{exp} > 0$. IDP sequences of test set with experimental data are provided in Supplementary Table S1.

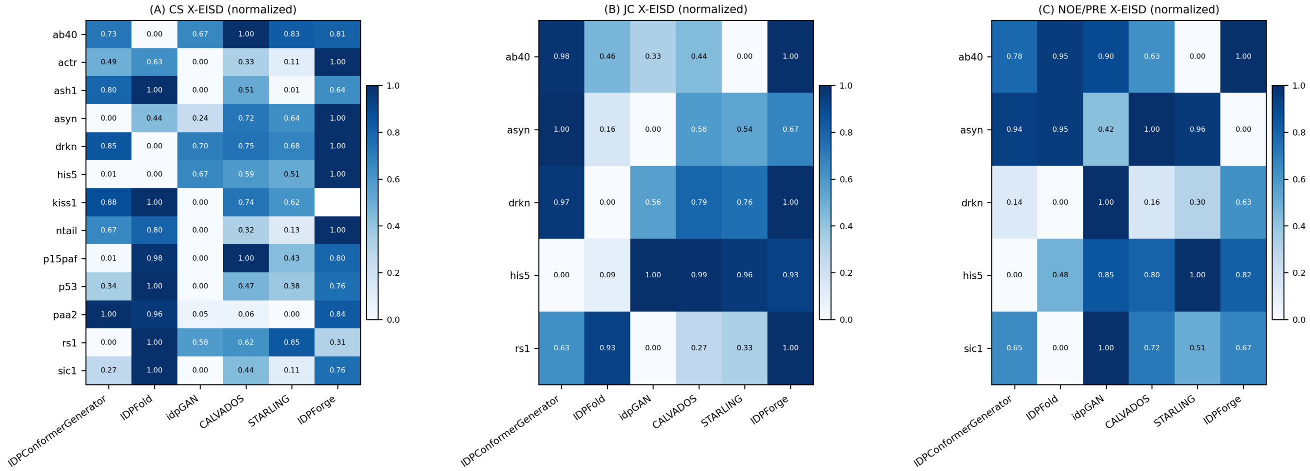


**Figure S4: Evaluation of the generated ensembles by experimental data type for the test sequences for various methods.** Aggregate X-EISD score normalized per protein for (A) chemical shifts, (B) J-Couplings, and (C) NOE and PRE data. X-EISD scores range between 0-1, and the higher score indicates better experimental agreement. IDP sequences of the test set with experimental data are provided in Supplementary Table S1.

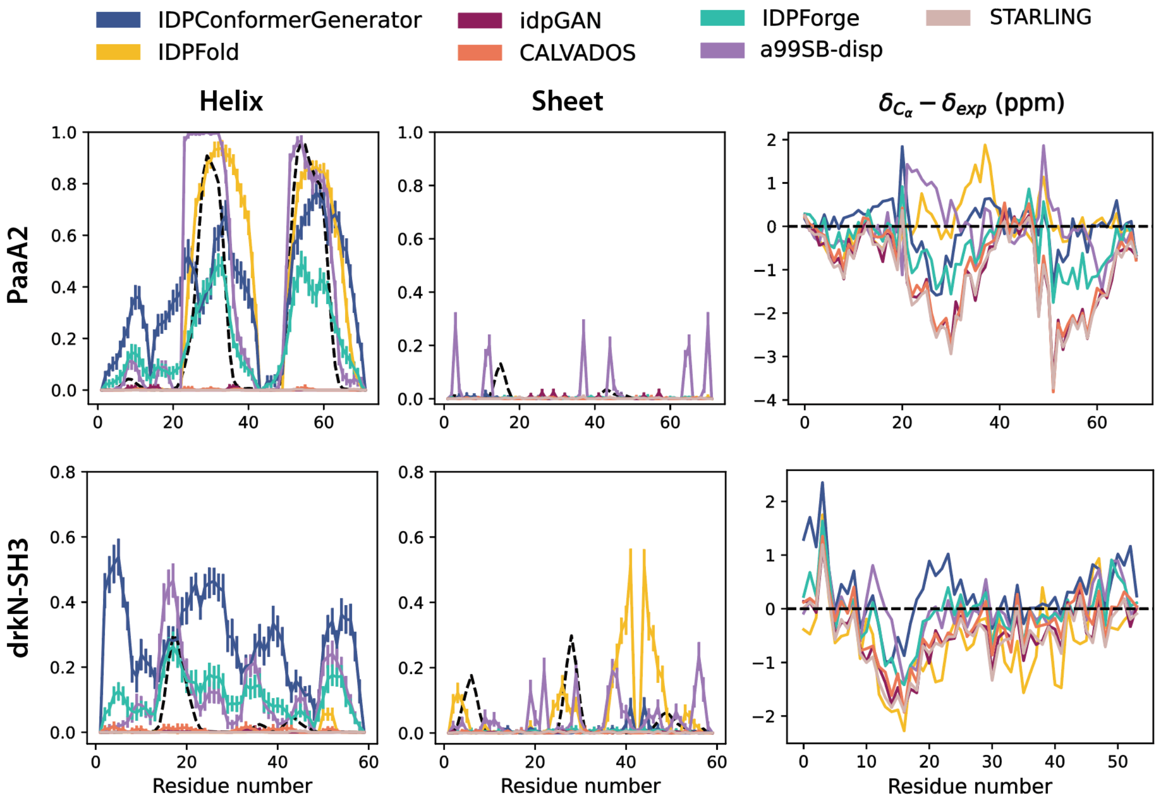


**Figure S5: DSSP propensity of the generated ensembles along with $C_\alpha$ chemical shift error for PaaA2 and drkN-SH3.** Left: DSSP propensity for helices and sheets. Error bars estimate the mean and standard deviation of 100 conformer ensembles from 30 trials. The black dashed line denotes the chemical shift inferred secondary structure from $\delta$2D. Right: $C_\alpha$ chemical shift error.

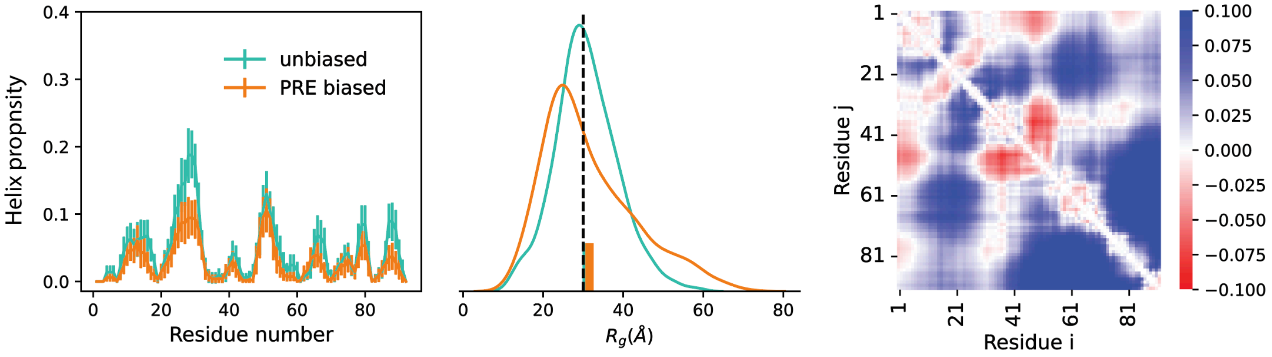


**Figure S6: Structural profiles for IDPForge unbiased Sic1 ensembles and biased with experimental data.** Left) DSSP propensity errorbars estimate the mean and standard deviation of 100 conformer ensembles from 30 trials. Center) $R_g$ profiles with experimental mean in black dashed line and ensemble means in bar plots. Right) Mean $C_\alpha$-$C_\alpha$ distance difference normalized by the IDPForge unbiased ensemble (calculated by $(\bar{d}_{PRE} - \bar{d}_{unbiased})/\bar{d}_{unbiased}$). Red indicates a compaction and blue for expansion.

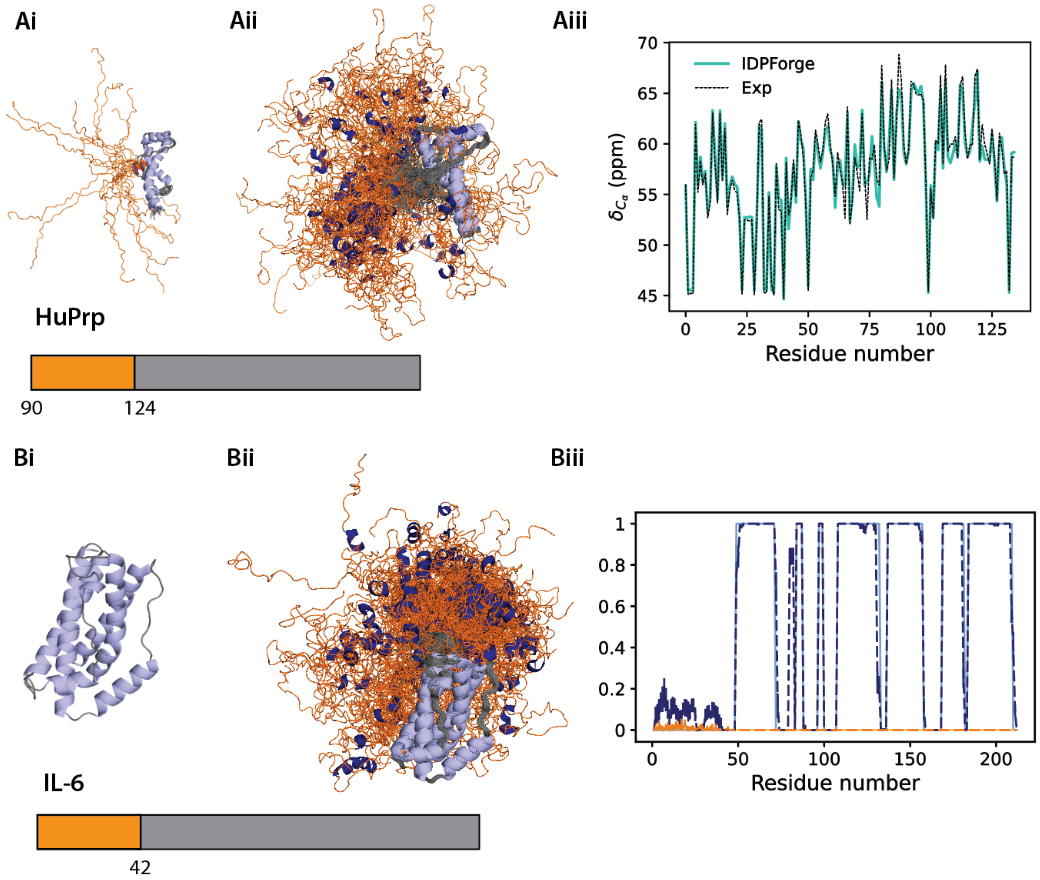


**Figure S7: All-atom IDPForge ensembles generated for IDRs with folded domains from experimental structures.** Ai) NMR ensemble HuPrp PDB ID 2LFT [73]. Bi) Interleukin-6 (IL-6) with 41 missing residues, PDB ID 4CNI chain C [74]. The generated IDPForge ensembles of (Aii) HuPrp and (Bii) IL-6 has folded regions in grey and IDRs in orange. (Aiii) $C_\alpha$ chemical shifts for IDPForge HuPrp ensemble and solution NMR [73]. (Biii) Secondary structure propensities for IL-6 of the generated ensembles in dashed lines and folded templates in solid and lighter colors. Error bars estimate the mean and standard deviation of 100 conformer ensembles from 30 trials.

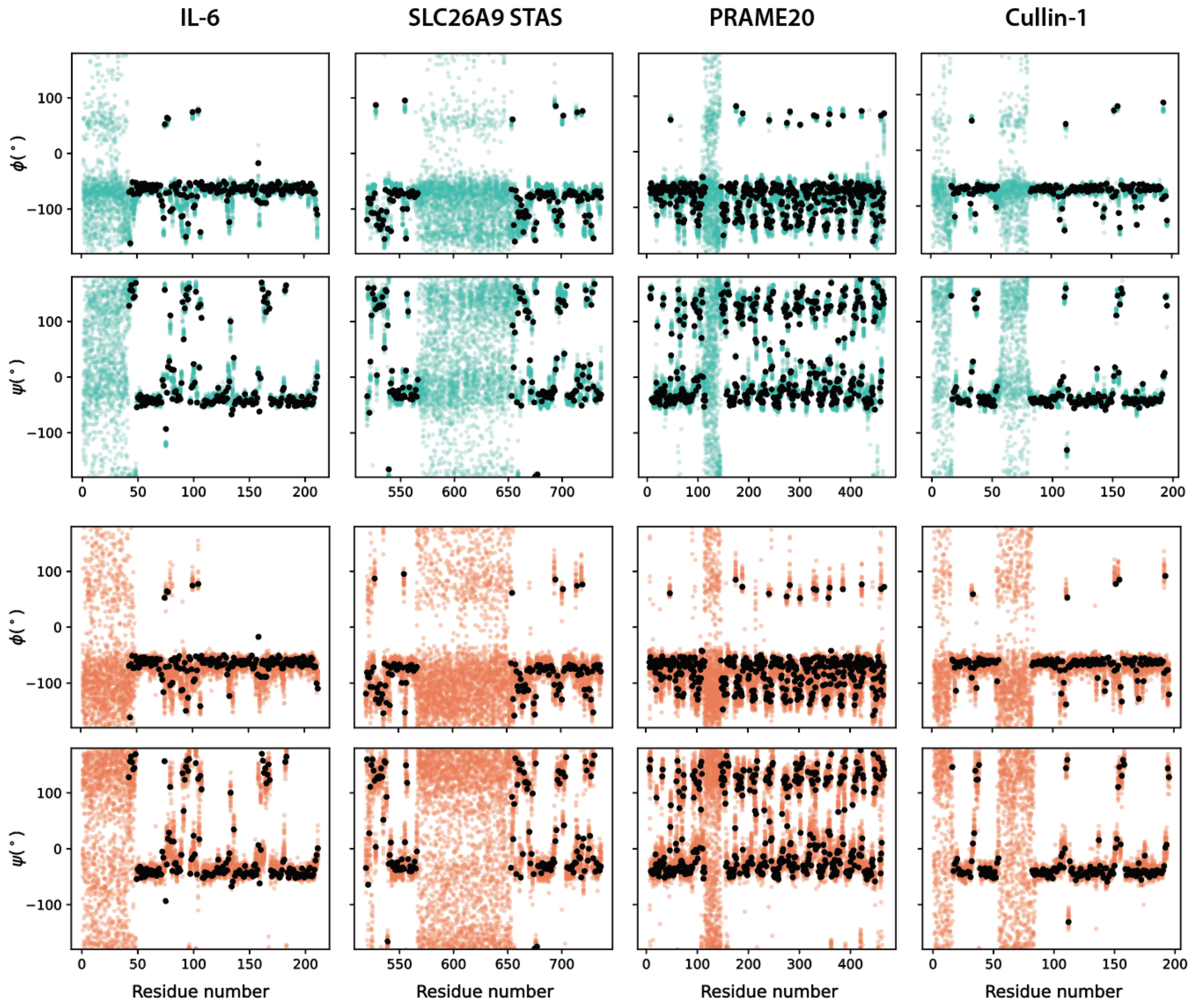


**Figure S8: Backbone scattering distributions for IL-6, SLC26A9 STAS, PRAME20 and Cullin-1 ensembles generated by IDPForge and CALVADOS.** Black dots denote the folded domain backbone torsions from each reference structure. Cyan: IDPForge; Orange: CALVADOS.

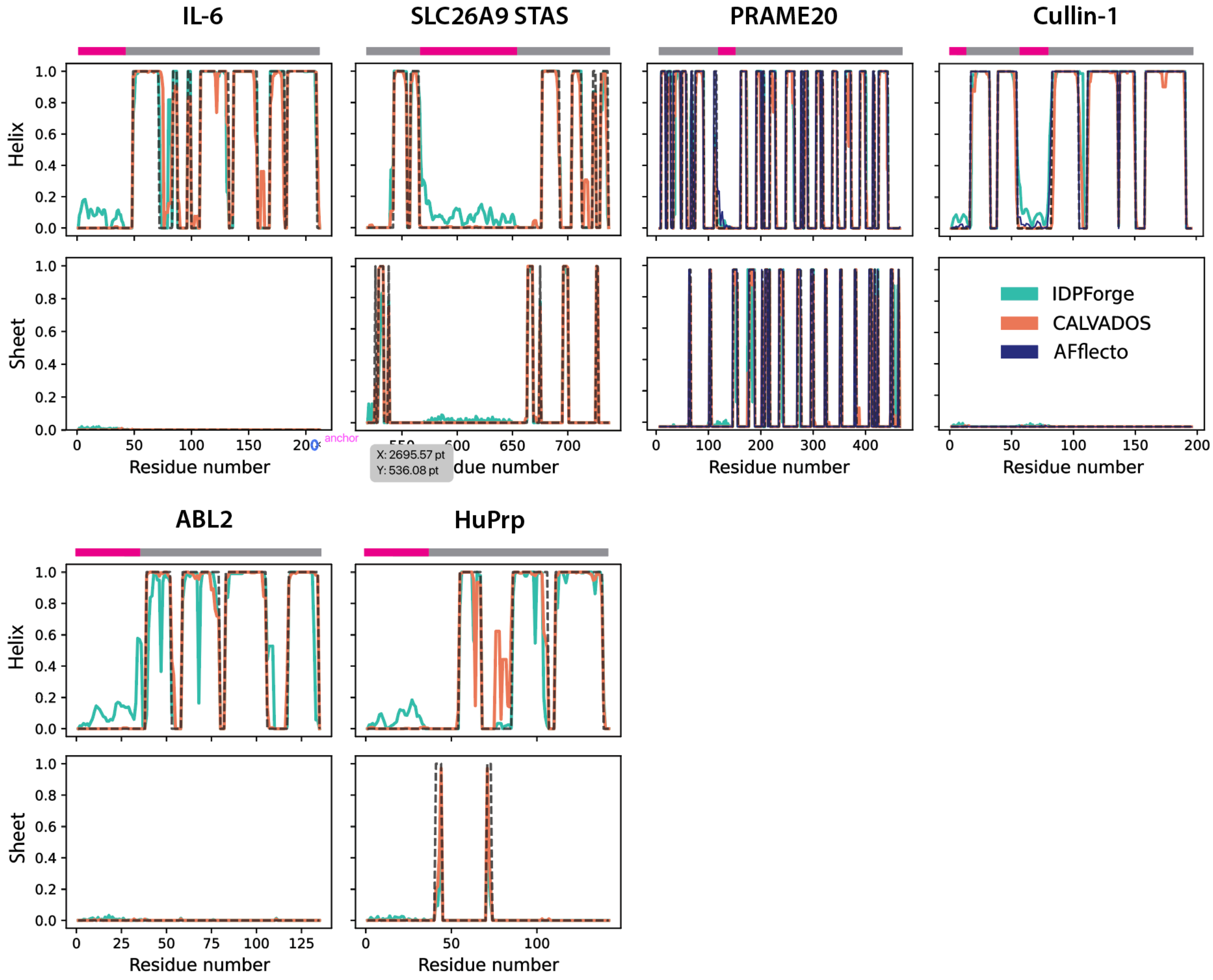


**Figure S9: DSSP propensity for ABL2, HuPrp, IL-6, SLC26A9 STAS, PRAME20 and Cullin-1 ensembles generated by IDPForge, CALVADOS and AFflecto.** DSSP propensities are averaged across ensembles of 100 conformers. The black dashed lines denote the folded domain DSSP calculated from each reference structure. Color bars on the top indicate the folded (grey) and disordered regions (magenta).

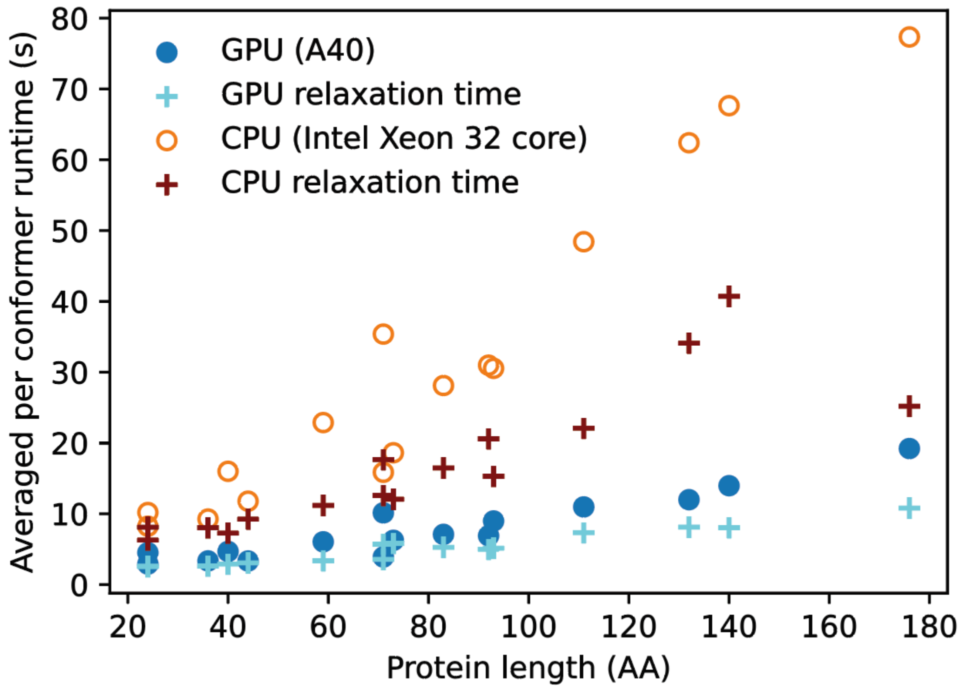


**Figure S10: Runtime scaling of IDPForge on CPU and GPU.**

[4] P Kingma Diederik. Adam: A method for stochastic optimization. *(No Title)*, 2014.

[5] Ting Chen, Ruixiang Zhang, and Geoffrey Hinton. Analog bits: Generating discrete data using diffusion models with self-conditioning. *arXiv preprint arXiv:2208.04202*, 2022.

[6] Joseph L Watson, David Juergens, Nathaniel R Bennett, Brian L Trippe, Jason Yim, Helen E Eisenach, Woody Ahern, Andrew J Borst, Robert J Ragotte, Lukas F Milles, et al. De novo design of protein structure and function with rfdiffusion. *Nature*, 620(7976):1089–1100, 2023.

[7] Adam Leach, Sebastian M Schmon, Matteo T Degiacomi, and Chris G Willcocks. Denoising diffusion probabilistic models on so (3) for rotational alignment. 2022.

[8] Valentin De Bortoli, Emile Mathieu, MJ Hutchinson, James Thornton, Yee Whye Teh, and Arnaud Doucet. Riemannian score-based generative modelling. *Advances in Neural Information Processing Systems*, page 46, 2022.

[9] Debashish Sahu, Monique Bastidas, and Scott A Showalter. Generating nmr chemical shift assignments of intrinsically disordered proteins using carbon-detected nmr methods. *Analytical biochemistry*, 449:17–25, 2014.

[10] Jakob Toudahl Nielsen and Frans AA Mulder. Chespi: chemical shift secondary structure population inference. *Journal of Biomolecular NMR*, 75(6):273–291, 2021.

[11] João MC Teixeira, Zi Hao Liu, Ashley Namini, Jie Li, Robert M Vernon, Mickaël Krzeminski, Alaa A Shamandy, Oufan Zhang, Mojtaba Haghighatlari, Lei Yu, et al. Idpconformergenerator: a flexible software suite for sampling the conformational space of disordered protein states. *The Journal of Physical Chemistry A*, 126(35):5985–6003, 2022.

[12] James Lincoff, Mojtaba Haghighatlari, Mickael Krzeminski, João M. C. Teixeira, Gregory-Neal W. Gomes, Claudiu C. Gradinaru, Julie D. Forman-Kay, and Teresa Head-Gordon. Extended experimental inferential structure determination method in determining the structural ensembles of disordered protein states. *Comm. Chem.*, 3(1):74, 2020. ISSN 2399-3669. doi: 10.1038/s42004-020-0323-0. URL `https://doi.org/10.1038/s42004-020-0323-0`.

[13] John Jumper, Richard Evans, Alexander Pritzel, Tim Green, Michael Figurnov, Olaf Ronneberger, Kathryn Tunyasuvunakool, Russ Bates, Augustin ídek, Anna Potapenko, Alex Bridgland, Clemens Meyer, Simon A. A. Kohl, Andrew J. Ballard, Andrew Cowie, Bernardino

Romera-Paredes, Stanislav Nikolov, Rishub Jain, Jonas Adler, Trevor Back, Stig Petersen, David Reiman, Ellen Clancy, Michal Zielinski, Martin Steinegger, Michalina Pacholska, Tamas Berghammer, Sebastian Bodenstein, David Silver, Oriol Vinyals, Andrew W. Senior, Koray Kavukcuoglu, Pushmeet Kohli, and Demis Hassabis. Highly accurate protein structure prediction with alphafold. *Nature*, 596(7873):583–589, 2021. ISSN 1476-4687. doi: 10.1038/s41586-021-03819-2. URL `https://doi.org/10.1038/s41586-021-03819-2`.

[14] Minkyung Baek, Frank DiMaio, Ivan Anishchenko, Justas Dauparas, Sergey Ovchinnikov, Gyu Rie Lee, Jue Wang, Qian Cong, Lisa N. Kinch, R. Dustin Schaeffer, Claudia Millán, Hahnbeom Park, Carson Adams, Caleb R. Glassman, Andy DeGiovanni, Jose H. Pereira, Andria V. Rodrigues, Alberdina A. van Dijk, Ana C. Ebrecht, Diederik J. Opperman, Theo Sagmeister, Christoph Buhlheller, Tea Pavkov-Keller, Manoj K. Rathinaswamy, Udit Dalwadi, Calvin K. Yip, John E. Burke, K. Christopher Garcia, Nick V. Grishin, Paul D. Adams, Randy J. Read, and David Baker. Accurate prediction of protein structures and interactions using a three-track neural network. *Science*, 373(6557):871–876, 2021. doi: 10.1126/science.abj8754. URL `https://doi.org/10.1126/science.abj8754`.

[15] Ruidong Wu, Fan Ding, Rui Wang, Rui Shen, Xiwen Zhang, Shitong Luo, Chenpeng Su, Zuofan Wu, Qi Xie, Bonnie Berger, et al. High-resolution de novo structure prediction from primary sequence. *BioRxiv*, pages 2022–07, 2022.

[16] Zeming Lin, Halil Akin, Roshan Rao, Brian Hie, Zhongkai Zhu, Wenting Lu, Nikita Smetanin, Robert Verkuil, Ori Kabeli, Yaniv Shmueli, et al. Evolutionary-scale prediction of atomic-level protein structure with a language model. *Science*, 379(6637):1123–1130, 2023.

[17] Namrata Anand and Tudor Achim. Protein structure and sequence generation with equivariant denoising diffusion probabilistic models. *arXiv preprint arXiv:2205.15019*, 2022.

[18] Bowen Jing, Bonnie Berger, and Tommi Jaakkola. Alphafold meets flow matching for generating protein ensembles. *arXiv preprint arXiv:2402.04845*, 2024.

[29] Mátyás Pajkos, Ilinka Clerc, Christophe Zanon, Pau Bernadó, and Juan Cortés. Afflecto: A web server to generate conformational ensembles of flexible proteins from alphafold models. *Journal of Molecular Biology*, page 169003, 2025.
